# Structure of human RNA Polymerase III

**DOI:** 10.1101/2020.06.29.177279

**Authors:** Ewan Phillip Ramsay, Guillermo Abascal-Palacios, Julia L. Daiß, Helen King, Jerome Gouge, Michael Pilsl, Fabienne Beuron, Edward Morris, Philip Gunkel, Christoph Engel, Alessandro Vannini

## Abstract

In eukaryotes, RNA Polymerase (Pol) III is the enzyme specialised for the transcription of the entire pool of tRNAs and several other short, essential, untranslated RNAs. Pol III is a critical determinant of cellular growth and lifespan across the eukaryotic kingdom. Upregulation of Pol III transcription is often observed in cancer cells and causative Pol III mutations have been described in patients affected by severe neurodevelopmental disorders and hypersensitivity to viral infection.

Harnessing CRISPR-Cas9 genome editing in HeLa cells, we isolated endogenous human Pol III and obtained a cryo-EM reconstruction at 4.0 Å. The structure of human Pol III allowed us to map the reported genetic mutations and rationalise them. Mutations causing neurodevelopmental defects cluster in hotspots that affect the stability and/or biogenesis of Pol III, thereby resulting in loss-of-function of the enzyme. Mutations affecting viral sensing are located in the periphery of the enzyme in proximity to DNA binding regions, suggesting an impairment of Pol III cytosolic viral DNA-sensing activity.

Furthermore, integrating x-ray crystallography and SAXS data, we describe the structure of the RPC5 C-terminal extension, which is absent in lower eukaryotes and not visible in our EM map. Surprisingly, experiments in living cells highlight a role for the RPC5 C-terminal extension in the correct assembly and stability of the human Pol III enzyme, thus suggesting an added layer of regulation during the biogenesis of Pol III in higher eukaryotes.

## INTRODUCTION

Transcription of the eukaryotic genome is mediated by three highly specialised nuclear RNA polymerase (Pol) enzymes. Pol III transcribes short untranslated RNAs which are essential for cellular functions, such as the entire pool of tRNAs, the precursor of the 5S ribosomal RNA and the U6 spliceosomal RNA^1^.

Pol III is a multi-subunit complex composed of 17 subunits. A central 10-subunit core which harbours the catalytic site and a peripheral heterodimeric stalk that are structurally conserved amongst the three eukaryotic Pols. The TFIIF-like RPC4/5 and the TFIIE-like RPC3/6/7 subcomplexes are Pol III specific and can be regarded as built-in general transcription factors that play a fundamental role in Pol III transcription initiation and termination^2-4^.

Across the eukaryotic kingdom, Pol III displays a high degree of conservation both in terms of subunit composition as well as sequence homology of the individual components. A notable exception is the subunit RPC5 which in metazoans encompasses a long C-terminal extension (RPC5EXT, approximately 450 residues long), whose function is currently unknown.

Pol III activity is highly regulated in a cell cycle and cell-type dependent manner^5^ and is a determinant of lifespan in eukaryotes^6^. In recent years, a large number of disease-causing mutations have been assigned to Pol III subunits, with a particular incidence of allele variants that strongly affect the correct development of the central nervous system, resulting in severe neurodegenerative diseases^7-14^. Furthermore, causative Pol III mutations have also been described in patients affected by hyper sensitivity to viral infection^15,16^.

To date, yeast Pol III has been extensively structurally and functionally characterised while its human counterpart has been left relatively untouched, due to the inherent technical challenges in obtaining yields amenable for structural biology. However, understanding the specific influence of pathological mutations and the role of regulatory elements unique to the human enzyme relies on such structural information. Here, we report the cryo-EM reconstruction of human Pol III. We further study the enzymes’ complete architecture using a structural biology hybrid approach integrating two crystal structures of the human RPC5 C-terminal extension as well as SAXS data and molecular modelling. Results of our comparative structural analysis rationalize the effect of pathological mutations and yield unexpected insights into Pol III regulation.

## RESULTS

### Purification of human RNA Polymerase III

In order to obtain a high-resolution structure of human Pol III we isolated the endogenous complex from HeLa cells. To this end, we employed CRISPR/Cas9 genome editing in human cells to create a homozygous knock-in of a cleavable GFP-tag at the C-terminus of subunit RPAC1 (shared between Pol I and Pol III) with (Figure 1a). Fractionation experiments followed by immuno-purification using an anti-GFP nanobody revealed that Pol III is present in both nuclear and cytoplasmic fractions (Figure 1b), in agreement with previous reports highlighting a Pol III cytosolic DNA-sensing activity^17,18^. An optimized large-scale purification, including an ion-exchange step to separate Pol I and Pol III, enabled the isolation of active human Pol III (Figure 1c,d; Extended Data Figure1) from total cell extracts with yields and quality amenable for further EM studies.

**Figure 1.**
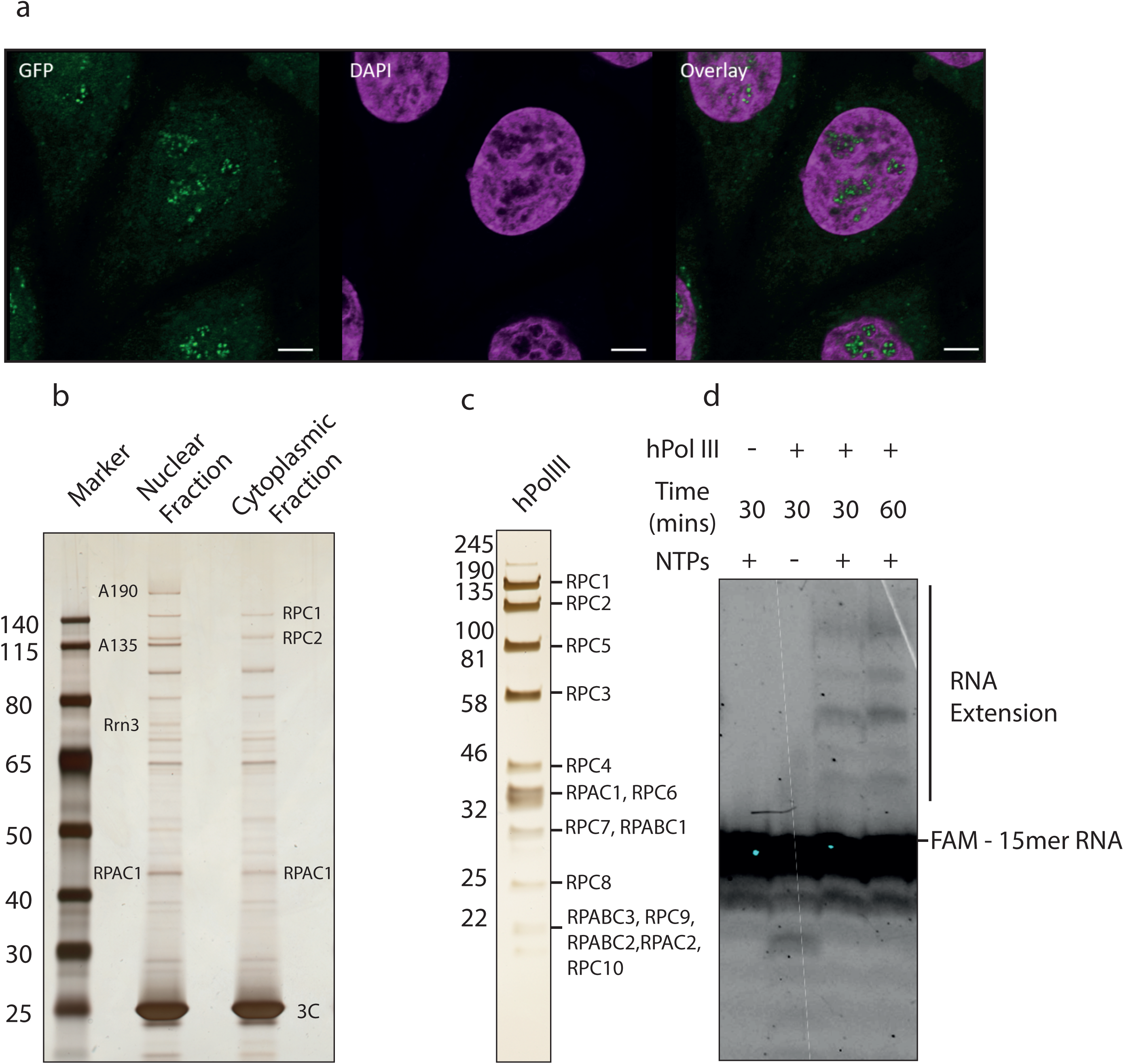
Purification of GFP-tagged endogenous human RNA polymerase III. **a**, Confocal imaging of modified HeLa cell line expressing homozygous sfGFP-tagged POLR1C gene. Endogenous GFP signal, representing Pol I and III (green), DAPI-staining (magenta), and overlay of both channels are shown. Scale bar: 5 µm. **b**, Affinity purified human RNA Pol III from HeLa nuclear and cytoplasmic fractions. Confirmed RNA Pol subunits are labelled. **c**, Purified human RNA Pol III (hPol III) from large-scale whole cell lysate with Pol III subunits marked. **d**, RNA extension assay of fluorescently-labelled FAM-15mer RNA primer by purified human Pol III. Marked is Pol III-mediated primer extension.

### Cryo-EM structure of human Pol III

The non-crosslinked purified human Pol III sample was applied to carbon-coated cryo-EM grids and imaged on a Titan Krios TEM microscope equipped with a Falcon III camera. Two data sets were collected at 0 and 30 degree tilting angles, to overcome preferred orientation of the sample on the cryo-grids, resulting in a merged dataset of 172,678 particles after 2D class averaging (Extended Data Figure 2, Table 1). The majority of imaged particles represented the intact 17-subunit human Pol III but a sizeable fraction with a similar angular distribution displayed no density for the RPC3/6/7 heterotrimer, which had possibly dissociated during purification, in agreement with earlier reports^19^, or during cryo-EM specimen preparation. Hierarchical 3D classification led to a reconstruction of the intact human Pol III from 25,369 particles at an overall resolution of 4.0 Å (Extended Data Figure 2-3; Table1). The core of the enzyme is characterised by a very detailed EM map where side chains are clearly discernible (Figure 2). The RPC8/9 stalk and the RPC3/6/7 subcomplex are more flexible than the core hence their local resolution is lower compared the rest of the complex (Extended Data Figure 3). Interestingly, the coiled-coil region of the clamp subdomain within the largest subunit RPC1, which is in direct contact with the RPC3/6/7 heterotrimer, also displays a high degree of flexibility. This finding suggests that the coiled-coil region of the clamp together with the heterotrimer form a discrete structural and functional unit, which in yeast has been shown to be able to sense melting of the upstream side of the transcription bubble^4^.

**Table 1.** Cryo-EM data collection, refinement and validation statistics

|  |  |
| --- | --- |
| **Data collection (0° Tilt)** |  |
| Voltage (kV) | 300 |
| Electron exposure (e<sup>-</sup>/Å<sup>2</sup>) | 44.1 |
| Defocus range (μm) | -1.0 to -3.0 |
| Pixel size (Å) | 1.065 |
| **Data collection (30° Tilt)** |  |
| Voltage (kV) | 300 |
| Electron exposure (e<sup>-</sup>/Å<sup>2</sup>) | 37.8, 40.6 |
| Defocus range (μm) | -1.2 to -3.0 |
| Pixel size (Å) | 1.065 |
| **Reconstruction (RELION)** |  |
| Initial particle images (no.) | 172,678 |
| Final particle images (no.) | 25,369 |
| Map resolution (Å) | 4.0 |
| FSC threshold | (0.143-thr) |
| Map sharpening *B* factor (Å<sup>2</sup>) | -116.839 |
| **Model composition** |  |
| Non-hydrogen atoms | 34636 |
| Protein residues | 4369 |
| **Refinement (PHENIX)** |  |
| Map CC | 0.53 |
| **R.m.s. deviations** |  |
| Bond lengths (Å) | 0.016 |
| Bond angles (°) | 1.086 |
| **Validation** |  |
| MolProbity score | 2.74 |
| Clashscore (all-atom) | 31.792 |
| Poor rotamers (%) | 0.0 |
| **Ramachandran plot** |  |
| Favored (%) | 78.66 |
| Allowed (%) | 20.94 |
| Disallowed (%) | 0.39 |

**Figure 2.**
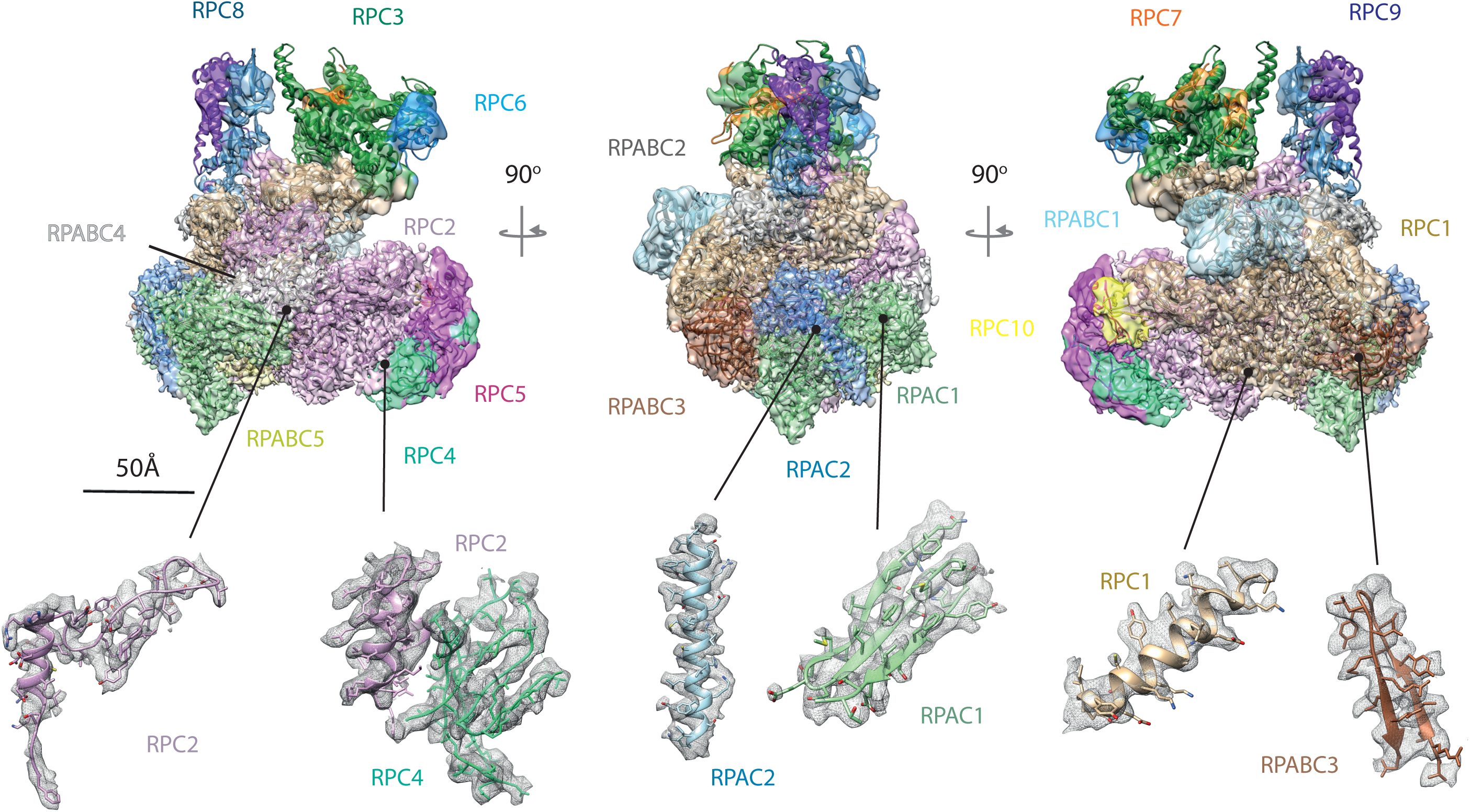
The structure of human RNA polymerase III. Shown is the electron density map filtered according to the local resolution with the fitted model shown in ribbon representation (above). Regions of the electron density map are coloured according to the subunit structure as labelled. Shown below are selected regions of several subunits showing the fit with the filtered electron density (mesh).

As can be expected from the high degree of sequence conservation, the overall structure of human Pol III resembles the yeast counterpart but also highlights local differences. In particular, two deletions in the human RPC1 subunit result in a rearranged, more compact foot domain (Extended Data Figure 4). Interestingly, similar structural rearrangements have been observed in the mammalian Pol II foot domain, a region that provides a binding interface for auxiliary regulators such as DSIF and the PAF complex^20,21^.

Due to the flexibility of peripheral subcomplexes, the reported RPC3 iron-sulphur cluster^22^ and the RPC5EXT, elements absent in the *S. cerevisiae* Pol III structures^3,4,23^, were not visible in our EM map.

### Structure of the RPC5 C-terminal extension

In order to gain insight into the structure and function of RPC5EXT, we determined the structure of its individual domains by x-ray crystallography (Figure 3; Table2). The RPC5EXT is formed by two consecutive tandem winged helix domains (tWHD1 residues 259-440; tWHD2 residues 556-708) connected by a 115 residue-long flexible linker. Such an architecture has not been reported for other components of the eukaryotic transcription apparatus and appears to be found exclusively in metazoan RPC5. Of the two tWHDs, tWHD1 is the most conserved while tWHD2 is absent in *C. elegans* and *D. melanogaster* (Extended Data Figure 5). The tWHD1 is formed by two juxtaposed winged helix domains that form a compact globular domain with one of the two recognition helices, typically involved in DNA binding, buried within the structure (Figure 3b). The compact conformation of tWHD1 is observed also in solution as highlighted by small-angle x-ray scattering (SAXS) data (Figure 3d, Extended Data Figure 6). The tWHD2 structure revealed a dimer formed by domain swapping (Figure 3c). This arrangement is likely caused by the crystallization conditions and, in agreement with this hypothesis, SAXS data showed a monomeric conformation as the most likely in solution (Figure 3e, Extended Data Figure 6). Nevertheless, the two possible conformations of tWHD2, compact or elongated, suggests a degree of flexibility within this domain. Finally, SAXS analysis of a construct encompassing the full-length RPC5EXT support the model of two globular compact tWHDs domains connected by a long flexible linker, spanning approximately up to 175Å in length (Figure 3f, Extended Data Figure 6-7).

**Table 2.** Rpc5-tWHD1 (aa. 259-440) Data collection, phasing and refinement statistics for MAD (SeMet) structures

|  | Crystal 1 (Native) |  | Crystal 2 (SeMet) |  |
| --- | --- | --- | --- | --- |
| **Data collection** |  |  |  |  |
| Space group | P6<sub>1</sub>22 | P6<sub>1</sub>22 | P6<sub>1</sub>22 | P6<sub>1</sub>22 |
| Cell dimensions |  |  |  |  |
| *a*, *b*, *c* (Å) | 56.30, 56.30, 275.88 | 56.43, 56.43, 277.27 | 56.44, 56.44, 277.17 | 56.41, 56.41, 276.95 |
| $\alpha$, $\beta$, $\gamma$ (°) | 90, 90, 120 | 90, 90, 120 | 90, 90, 120 | 90, 90, 120 |
|  |  | Peak | Inflection | Remote |
| Wavelength | 0.91983 | 0.97965 | 0.97980 | 0.971970 |
| Resolution (Å) | 48.01 - 2.23 (2.29 - 2.23) | 48.87 - 2.72 (2.79 - 2.72) | 48.88 - 2.63 (2.70 - 2.63) | 48.11 - 2.50 (2.56 - 2.50) |
| *R*<sub>sym</sub> or *R*<sub>merge</sub> | 0.05 (0.749) | 0.324 (2.163) | 0.268 (1.820) | 0.246 (1.656) |
| *I* / σ*I* | 26.9 (5.7) | 8.7 (1.5) | 9.4 (1.5) | 8.7 (1.2) |
| Completeness (%) | 100.0 (100.0) | 100.0 (100.0) | 100.0 (100.0) | 100.0 (100.0) |
| Redundancy | 27.4 (28.8) | 21.6 (19.6) | 21.2 (15.8) | 20.0 (10.6) |
| **Refinement** |  |  |  |  |
| Resolution (Å) | 45.97 - 2.23 (2.31 - 2.23) |  |  |  |
| No. reflections | 376023 |  |  |  |
| *R*<sub>work</sub> / *R*<sub>free</sub> | 0.1800 / 0.2193 |  |  |  |
| No. atoms | 1496 |  |  |  |
| Protein | 1371 |  |  |  |
| Ligand/ion | 4 |  |  |  |
| Water | 121 |  |  |  |
| *B*-factors |  |  |  |  |
| Protein | 50.52 |  |  |  |
| Ligand/ion | 89.50 |  |  |  |
| Water | 54.85 |  |  |  |
| R.m.s deviations |  |  |  |  |
| Bond lengths (Å) | 0.008 |  |  |  |
| Bond angles (°) | 0.84 |  |  |  |

**Table 2 Rpc5-tWHD2 (aa. 556-708)** Data collection, phasing and refinement statistics for MAD (SeMet) structures
|  | Crystal 1 (SeMet) |  | Crystal 2 (SeMet) |  |
| --- | --- | --- | --- | --- |
| **Data collection** |  |  |  |  |
| Space group | P1 2<sub>1</sub> 1 | P1 2<sub>1</sub> 1 | P1 2<sub>1</sub> 1 | P1 2<sub>1</sub> 1 |
| Cell dimensions |  |  |  |  |
| *a*, *b*, *c* (Å) | 41.11, 75.76, 62.4 | 41.37, 75.67, 62.77 | 41.35, 75.78, 62.79 | 41.30, 75.72, 62.67 |
| $\alpha$, $\beta$, $\gamma$ (°) | 90.0, 103.57, 90.0 | 90.0, 103.73, 90.0 | 90.0, 103.72, 90.0 | 90.0, 103.64, 90.0 |
| Wavelength | 0.97942 | Peak<br>0.97942 | Inflection<br>0.97957 | Remote<br>0.97174 |
| Resolution (Å) | 30.33 - 1.48 (1.52 - 1.48) | 47.48 - 1.74 (1.79 - 1.74) | 47.52 - 1.71 (1.75 - 1.71) | 60.90 - 1.78 (1.83 - 1.78) |
| *R*<sub>sym</sub> or *R*<sub>merge</sub> | 0.075 (0.735) | 0.095 (0.942) | 0.088 (0.954) | 0.118 (1.001) |
| *I* / $\sigma I$ | 11.8 (1.3) | 10.5 (1.6) | 11.2 (1.5) | 9.0 (1.5) |
| Completeness (%) | 96.1 (71.2) | 99.9 (100.0) | 99.9 (100.0) | 99.9 (100.0) |
| Redundancy | 6.1 (3.6) | 6.7 (5.8) | 6.7 (5.4) | 6.8 (6.4) |
| **Refinement** |  |  |  |  |
| Resolution (Å) | 30.33 - 1.55 (1.60 - 1.55) |  |  |  |
| No. reflections | 364951 |  |  |  |
| *R*<sub>work</sub> / *R*<sub>free</sub> | 0.1786 / 0.2078 |  |  |  |
| No. atoms | 3053 |  |  |  |
| Protein | 2527 |  |  |  |
| Ligand/ion | 8 |  |  |  |
| Water | 518 |  |  |  |
| *B*-factors |  |  |  |  |
| Protein | 24.28 |  |  |  |
| Ligand/ion | 20.98 |  |  |  |
| Water | 36.33 |  |  |  |
| R.m.s deviations |  |  |  |  |
| Bond lengths (Å) | 0.011 |  |  |  |
| Bond angles (°) | 1.15 |  |  |  |

**Figure 3.**
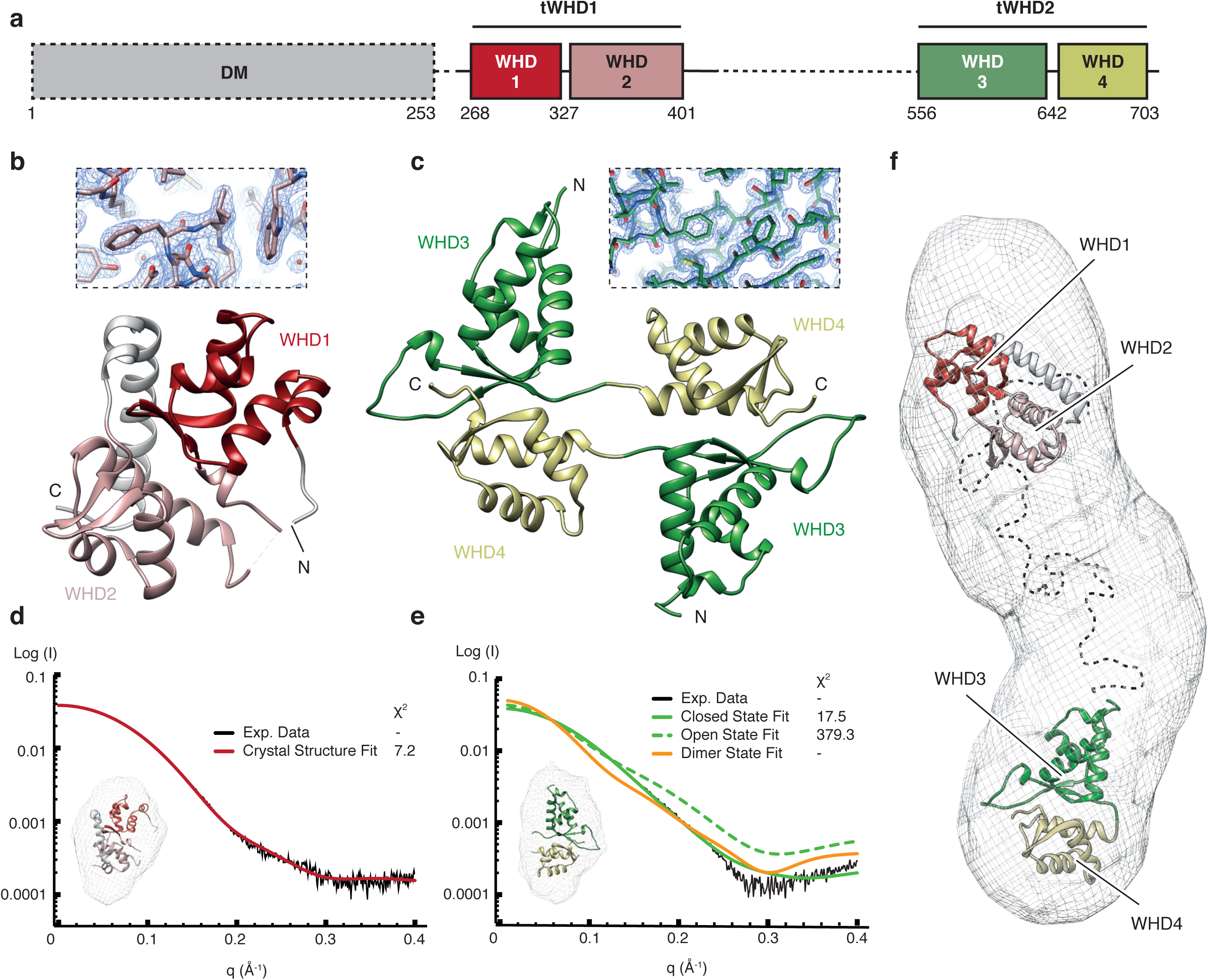
Structure of the Rpc5 subunit C-terminal region. **a**, Domain architecture of Rpc5 C-terminus. tWHD1 and tWHD2 are coloured in red and green shades, respectively. Regions are depicted according to their presence (black line) or absence (black dashed line) in the crystal structures. DM refers to Rpc5 dimerization module. **b**, Crystallographic model of Rpc5-WHD1 (dark red), Rpc5-WHD2 (light red) and the linker regions (grey) in ribbon cartoon. Inset shows detail of the electron density map. **c**, Overall structure of Rpc5-tWHD2 crystallographic packing in ribbon cartoon. Rpc5-WHD3 and Rpc5-WHD4 are shown in dark and light green, respectively, and the inset shows detail of the electron density map. **d**, Fitting of Rpc5-tWH1 (red, inset) into the SAXS experimental data (black). **e**, Fitting of Rpc5-tWH2 in “closed” (green, inset), “open” (dashed green) and dimer (orange) conformations into the SAXS experimental data (black). **f**, Docking of Rpc5-tWH1 (red) and Rpc5-tWH2 (green) into the ab initio SAXS envelope generated from Rpc5EXT SAXS data collection.

Comparison with existing protein structures using the Dali server (http://ekhidna.biocenter.helsinki.fi/dali_server/) suggested similarities between tWHD1 and the WHD of S. cerevisiae Pol II general transcription factor TFIIF Rap30 subunit^24-26^ and with the tWHD of Pol I A49 subunit^27-29^. Both subunits are orthologs of RPC5 and involved in stabilization of the pre-initiation complexes (PICs), suggesting a putative functional link. However, while the position of TFIIF Rap30 WHD in the Pol II PIC clashes with the Bdp1 subunit of transcription factor TFIIIB in the Pol III PIC^3,4^ (Figure 4), the equivalent position of A49 tWHD in the Pol I PIC^27^ is accessible in the Pol III PIC (Figure 4). Thus, one possibility is that, analogously to A49 tWHD, the RPC5 tWHD1 participates in an interaction with the upstream DNA and bound transcription factors, thus stabilising the human Pol III PIC. Additionally, the Dali server analysis retrieved similarities between the individual WHDs of RPC5EXT tWHD2 and the WHDs of cullin and cullin-like proteins, which are involved in ubiquitin-dependent proteolysis^30^.

**Figure 4.**
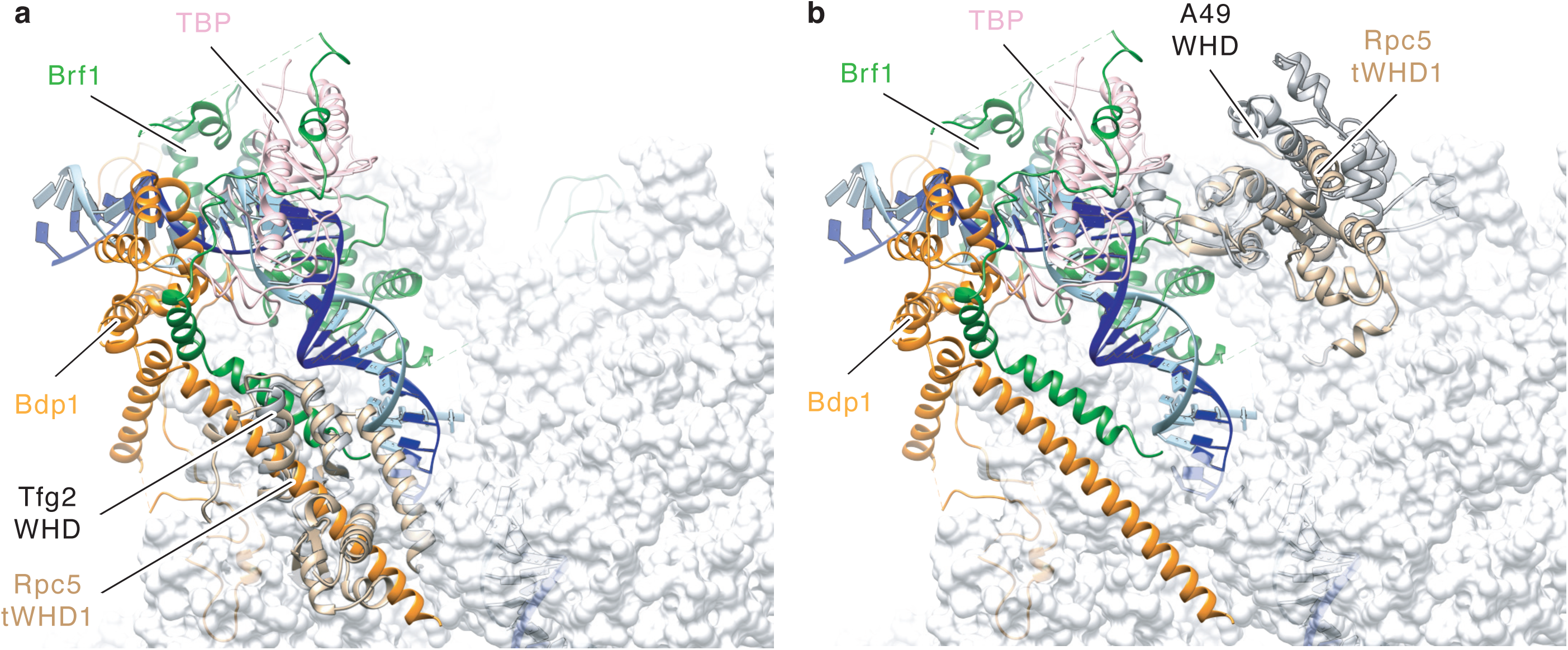
Structural alignment of RPC5 tWHD1 to Pol I and Pol III PICs. **a**, Structural alignment between S. cerevisiae RNA Pol II (PDB code: 5FYW) and RNA Pol III (PDB code: 6EU0). Rpc5 tWHD1 (wheat) superimposition to yeast Tfg2 WHD (gray) leads to clashes with Bdp1 subunit stem motif (orange). **b**, Structural alignment between S. cerevisiae RNA Pol I (PDB code: 5W66) and RNA Pol III (PDB code: 6EU0). Rpc5 tWHD1 (wheat) superimposition to yeast A49 tWHD (gray) does not cause clashes with TFIIIB subunits Brf1 (green), TBP (pink) or Bdp1 (orange).

### The RPC5 extension is required for RNA Pol III stability

In order to gain insight into the functional role of RPC5EXT, we used siRNA to knock down RPC5 in HEK293T cells and rescued it with ectopic expression of HA-tagged RPC5 constructs encompassing the full length protein (RPC5FL) or a version of RPC5 devoid of the RPC5EXT (RPC5ΔC) (Extended Data Figure 8). Immunoprecipitation using anti-HA magnetic beads revealed that both RPC5FL and RPC5ΔC are able to integrate into and pull-down a bona fide intact Pol III complex, as probed by RPC1, RPC2 and RPC4 antibodies (Extended Data Figure 8b). However, the corresponding immunoblots of whole cell-extracts, prior to the immunoprecipitation, indicate lower steady-state levels of RPC5ΔC compared to RPC5FL, pointing towards a direct role of RPC5EXT in enhancing RPC5 stability. In order to further explore the role of RPC5EXT in regulating RPC5 stability in the context of an intact Pol III complex, we employed a cycloheximide chase assay (Figure 5). Levels of RPC5FL remained stable for the course of the experiment (8 hours), as well as subunits RPC1 and RPC2, suggesting a relatively long half-life of the Pol III complex (Figure 5a). On the contrary, RPC5ΔC was rapidly degraded and almost completely depleted after only 2 hours following cycloheximide treatment (Figure 5b). Surprisingly, subunits RPC2 and, to a minor extent, RPC1 were also rapidly depleted, suggesting that RPC5EXT is essential for the stability of the whole Pol III complex.

**Figure 5.**
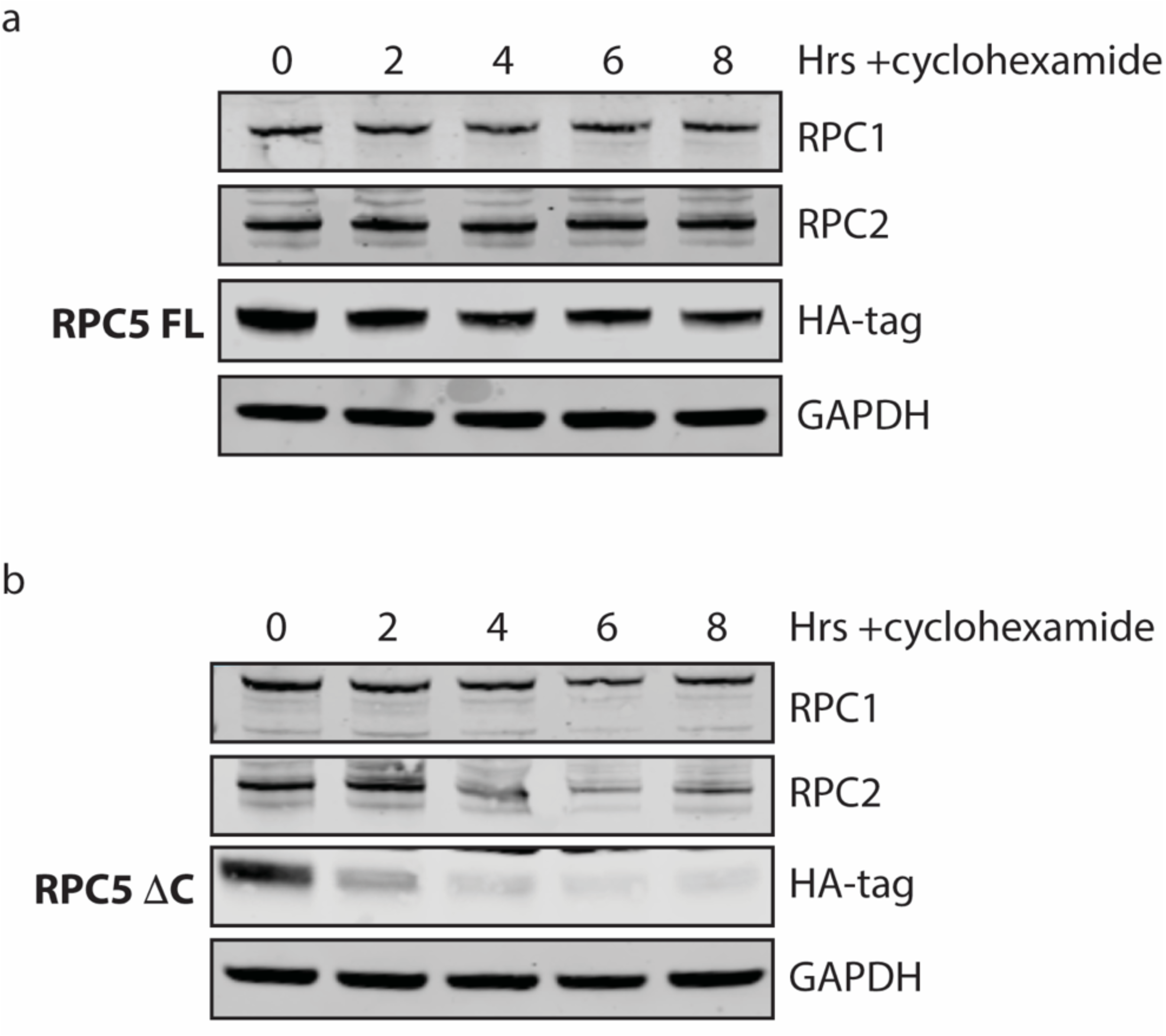
Cycloheximide chase assay investigation of RPC5 protein stability. HEK293T cells were seeded, endogenous RPC5 was knocked down via siRNA and either the FL (**a**) or ΔC (**b**) RPC5 constructs were transfected. After 24 hours, cycloheximide was added in at a concentration of 300*µ*g/ml and cells lysed at the specified time points. RPC1, 2 and 5 (as shown via HA-tag antibody) levels were probed by western blot.

### Pathologic genetic mutations map to Pol III subunit interfaces

Many studies have reported mutations of the Pol III enzyme that are related to human diseases, in particular heritable diseases which affect the correct development of the central nervous system (CNS). Specifically, allele variants encoding mutated versions of the Pol III subunits RPC1, RPC2, RPAC1 and RPAC2 subunits have been established as causative mutations of hypomyelinating leukodystrophy (HL)^7-10,31-35^, Treacher-Collins syndrome (TCS)^11,12^ and Wiedemann-Rautenstrauch syndrome (WRS)^13,14^.

To rationalize these findings, we mapped known Pol III mutations on our high-resolution structure (Figure 6, Extended Data Table 1). Reported mutation affecting CNS development tend to cluster in specific hotspots, very often at the interface of several Pol III subunits. For example, TCS mutations L51R and T50I in RPAC2 result in disruption of hydrophobic and salt-bridge interactions, respectively, at the interface with the RPAC1 subunit, suggesting a strong destabilising effect that might impair correct assembly of the enzyme (Figure 6). Analogously, most reported HL mutations lay at the interface of several subunits and have disruptive effects on these interfaces (Figure 6, Extended Data Table 1). Interestingly, WRS mutation R1069Q in subunit RPC1 disrupts a charged interaction with residue N1249 in the same subunit. This residue is itself mutated in HL, possibly altering the interface between subunits RPC1 and RPABC1 in both pathologies. Overall, these finding indicate a general molecular mechanism from mutations resulting in CNS disorders, which is the partial loss of function of Pol III activity through destabilisation of the enzyme core.

**Figure 6.**
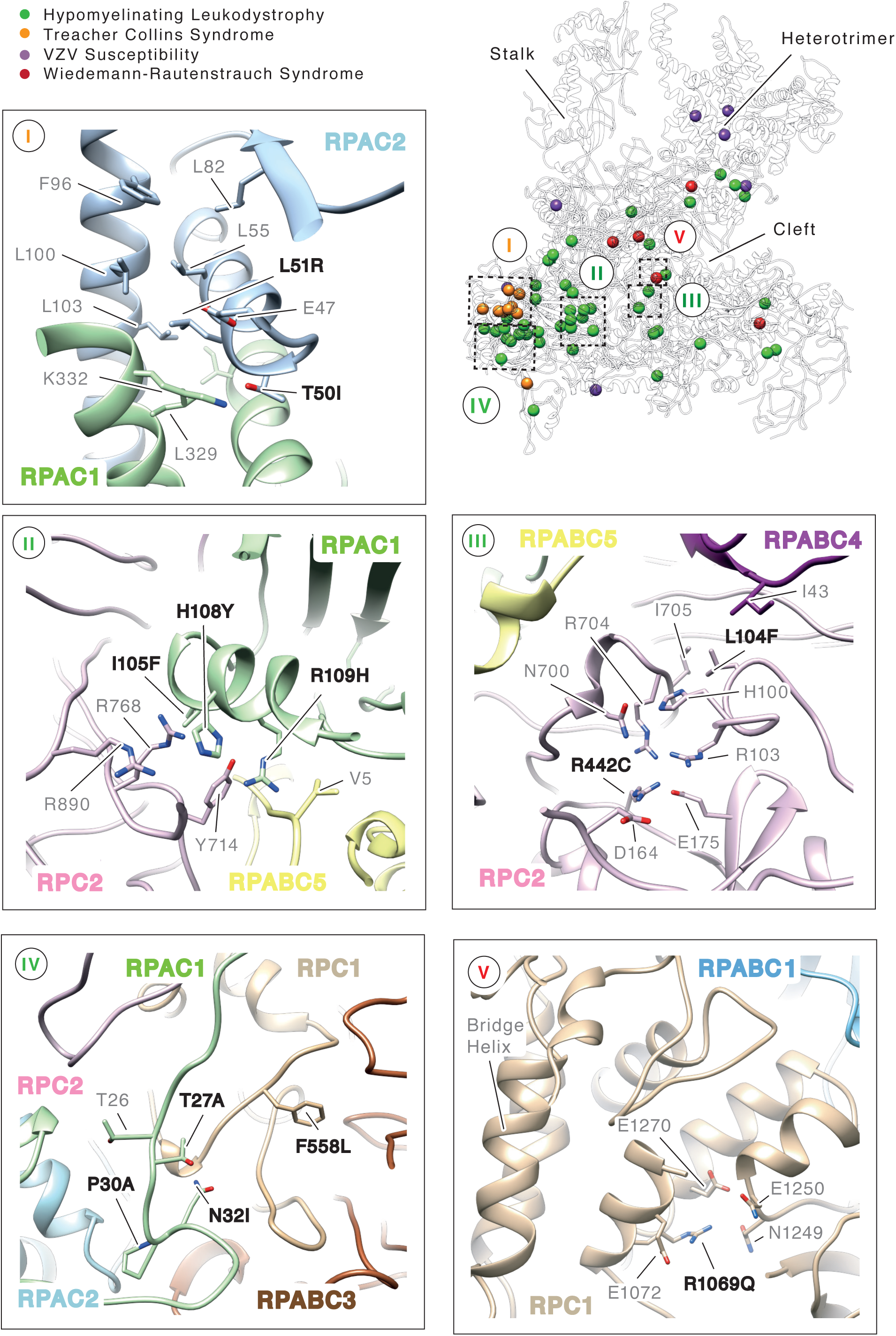
Mapping genetic mutations in human RNA Polymerase III. The position of residues mutated in hypomyelinating leukodystrophy (HL, green), Treacher-Collins syndrome (TCS, orange), Wiedemann-Rautenstrauch syndrome (WRS, red) and VZV susceptibility (VZV, purple) is shown as solid spheres. The close-up panels show details of the interfaces involved in TCS (panel I), HL (panels II-IV) and WRS mutations (panel V). Side chains are shown in stick representation and Pol III subunits are coloured as in Figure 2.

Recently, Pol III mutations have also been described in patients affected by acute severe response to Varicella zoster virus (VZV) infection^15,16,36^. Most of these mutations map at the periphery of the Pol III enzyme and result in neutralising basic charged residues exposed to the solvent in proximity of DNA binding regions (Figure 6). Since human Pol III has been shown to display cytosolic DNA-sensing activity, it is conceivable that VZV mutations indeed impair proper DNA binding and transcription factor-independent RNA synthesis in the cytoplasm.

Overall, these findings are in agreement with previous work using the homologous yeast Pol III and/or Pol I and Pol II enzymes to map disease-causing mutations^8-10,15,37^. However, the availability of a high-resolution structure of human Pol III enable the comprehensive mapping and rationalisation of these allele variants with high confidence.

## DISCUSSION

Here we describe the 4.0 Å resolution cryo-EM structure of apo human Pol III. The structure confirms the overall high-degree of structural homology with its *S. cerevisiae* counterpart but also highlights specific differences, such as a rearranged foot domain (Extended Data Figure 4). Integrating cryo-EM data with x-ray crystallographic and SAXS data for the metazoan-specific RPC5EXT, detailed structural information have been obtained for the whole Pol III complex (Extended Data Figure 9). Surprisingly, experiments in living cells highlighted a prominent role of RPC5EXT for the integrity of the Pol III complex. Absent in lower eukaryotes, RPC5EXT thus represents an additional metazoan-specific module, impacting on the correct assembly of Pol III. Several abundant phosphorylation sites have been identified in RPC5EXT which, together with the evidence of structural similarities between RPC5EXT tWHD2 and factors involved in targeted degradation, suggests the intriguing hypothesis of an RPC5EXT-mediated layer of regulation impacting overall Pol III abundance in response to environmental cues that may have evolved in higher organisms.

Furthermore, the high-resolution structure of human Pol III enabled the mapping of more than 85% of reported Pol III genetic mutations with high precision, rationalising their effects at a molecular level (Extended Data Table 1). Mutations affecting the CNS development tend to spatially cluster together and it seems likely that the severity of the phenotypes observed in HL, TCS and WRS correlate with the disruptive effect of such variants. For example, TCS mutations appear to be particularly disruptive at the interface of RPAC1 and RPAC2, two subunits shared between Pol I and Pol III. Interestingly, mutations N32I and N74S in RPAC1 mutations associated with HL lead to reduced Pol III assembly and nuclear import, without affecting Pol I^7^. This is consistent with our model as these residues mediate interactions with Pol III-specific RPC1 and RPC2, respectively (Extended Data Table 1).

Finally, since Pol III represents a central nexus involved in the regulation of organismal growth, development and life span in eukaryotes and is often deregulated in cancer, the structure of the human enzyme will represent an invaluable tool to aid the design of small molecules capable of specifically targeting Pol III transcription for therapeutic purposes.

## MATERIALS AND METHODS

### CRISPR/Cas9 genome editing

HeLa cells were cultured in DMEM medium (D6429, Sigma Aldrich) supplemented with 10% FBS (P40-37500, PAN-Biotech) and 1% AAS (A5955, Sigma Aldrich). Genomic integration of sfGFP ORF at the C-terminus of POLR1C was done by CRISPR/Cas9 according to a published protocol (Ran *et al*, 2013b) with some modifications.

Design of the guide RNAs (gRNAs) was done with a web-based tool (http:\CRISPR.mit.edu) and annealed oligos (gRNA1 = gCTAGTTCATCCAAGAAGCGC; gRNA2 = gCGGTTCAGATGGACTGAGCT) were cloned via BplI into the bicistronic Cas9n expression vector pSpCas9n(BB)-2A-Puro (PX462) V2.0, which was a gift from Feng Zhang (Addgene plasmid #62987; http://n2t.net/addgene:62987; RRID: Addgene_62987). A donor plasmid carried a short GS-linker sequence with an embedded HRV 3C protease cleavage site and the sfGFP ORF surrounded by two large sequence segments homologous to the insertion locus in the genome.

HeLa cells were transfected with a 1:1:1 mix of gRNA1 and gRNA2 vectors together with the donor plasmid using PolyJet transfection reagent (SL100688, SignaGen Laboratories) according to the manufacturer’s instructions. Several days later the GFP-expressing cells were enriched by flow cytometry using a Bio-Rad S3e cell sorter. GFP-positive cells were seeded on large culture dishes such that they could grow as single cell colonies. After 2-3 weeks, colonies were transferred manually into multi-well slides for live cell imaging and were screened under identical microscope settings. The brightest clones were selected for expansion. These monoclonal populations were validated by PCR on extracted genomic DNA (using the Blood & Tissue Kit, Qiagen).

The selected cell line was cultivated adherently and adapted to suspension growth as follows: Cells from 8 flasks (about 70×10^6^ cells total; 83.3912.302, Sarstedt) were detached by incubation with trypsin (25300, Gibco) at 37°C for 5 min, transferred to a spinner flask (250 mL total volume; 4500, Corning) and cultured in suspension with high-glucose DMEM (11965, Gibco) supplemented with 1% FBS (10270, Gibco) and 1% Penicillin/Streptomycin (P0781, Sigma Aldrich) under moderate stirring at 37°C, 5% CO_2_ atmosphere. To expand the culture, 1x the current volume of fresh media including all supplements was added when cells reached a density of ∼3,5×10^5^ cells/mL, and transferred to spinner flasks of increasing volume when required. Cells were harvested by centrifugation and washed with PBS before flash-freezing the pellet.

For fluorescence imaging, cells were grown adherently on cover slips to 50% confluency. After washing the cells with pre-warmed (37°C) PBS, they were fixed with 3.7% paraformaldehyde in PBS for 10 minutes at 37°C. The fixation was stopped by addition of 100 mM glycine in PBS for 5 min at 37°C and cells were washed twice with PBS. The cells on the cover slips were mounted on the specimen slide with the help of a drop Prolong Gold Antifade Mountant with DAPI (P36941, Thermo Fisher Scientific) and dried for at least three days in the dark.

The fluorescent specimens were imaged using a Zeiss Axio Observer.Z1 / 7 microscope and a 63x oil objective lens. DAPI staining was detected with the help of a 405 nm excitation laser and for the emission a wide band pass filter (300-720 nm) was used. For sfGFP detection a 488 nm laser and the same band pass filter (300-720 nm) was applied. The images were captured with the Airyscan mode and detector. Processing was done using the Zeiss AxioVision software, the Zeiss ZEN 3.0 (ZEN lite) software and Fiji.

### Anti-GFP pulldown from nuclear and cytosolic fractions

A total of ∼7L POLR1C-GFP Hela cells were grown in a spinner flask (Corning) and harvested by centrifugation, yielding a cell pellet of 8.1g (estimated total of 2.3×10^9 cells). Nuclear extracts were prepared as described^38^. This resulted in a nuclear pellet of 5.1g, corresponding to 12 mL of nuclear extract in ‘Roeder C Buffer’ (25% v/v glycerol, 20mM HEPES-KPOH pH 7.9, 1.5 mM MgCl_2_, 0.2 mM EDTA pH 8.0, 420 mM NaCl, 0.5 mM DTT, 0.5 mM PMSF). Nuclear extracts and cytosolic fraction were split into fractions of 0.5 mL each.

An aliquot of nuclear extract and cytosolic fraction were each supplemented with 500 *µ*L wash buffer (5% v/v glycerol, 20 mM HEPES 7.8, 420 mM NaCl, 1 mM MgCl_2_, 0.1 mM ZnCl, 2 mM beta mercapto-ethanol, 0.5 mM PMSF) and 5 units DNase I (Thermo Fished Scientific) followed by 30 min incubation at 4°C in an overhead mixer. Debris was removed by centrifugation (Eppendorf Centrifuge 5427 R, Rotor FA-45-12-17) 13k rpm, 4°C, 30 min. Nuclear extracts and cytosolic fraction were each bound to 20 *µ*L (slurry) ‘GFP-selector’ Beads (NanoTag) for 2 hours and washed twice with 0.5 mL wash buffer times and washed/eluted as described above. Elution was carried out with 3C protease in 50 *µ*L wash buffer at 4°C for 2h. Pre-cast SDS-PAGE gels (4-12% NuPAGE Bis-Tris, Thermo Scientific) were loaded with 12.5 *µ*L fractions.

### Large-Scale Human RNA Polymerase Purification

Large-scale cell growth was carried out at the Cell Services Scientific Technical Platform at The Francis Crick Institute, London. Adherent HeLa POLR1C-GFP cells were grown in DMEM-4 medium supplemented with 1% FCS, 1% Glutamax and 1% Penicillin/Streptomycin. Confluent cells were harvested by trypsin treatment followed by gentle centrifugation. The cell pellet was subsequently resuspended in RPMI-1640 supplemented with 5% FCS, 1% Glutamax and 1% penicillin/streptomycin to allow for cell growth in suspension. Cells were expanded in suspension using a small glass spinner flask flushed with CO_2_ at 37°C. Cells were expanded to a maximum volume of 1.2L per growth in a 3L glass spinner flask. Cells were grown to a density of to 1×10^6^ cells/ml with viability maintained at greater than 90%. Following growth, cells were collected by gentle centrifugation at room temperature. The resulting cell pellets were washed with PBS and cells pelleted again via centrifugation. The final cell pellets were stored at -80°C prior to purification.

For large scale purification of human RNA polymerase, whole cell lysate was produced from a cell pellet derived from 20L of HeLa cells grown to 1×10^6^ cells/ml density. The cell pellet was resuspended in lysis buffer (50mM Tris HCl pH 8.0, 250mM (NH_4_)_2_SO_4_, 20% v/v glycerol, 1mM MgCl_2_, 10μM ZnCl_2_, 10mM β-mercaptoethanol) and two protease inhibitor tablets (Roche) added. Lysis was performed through repeated passage of the cell suspension through a dounce followed by sonication with the resulting lysate cleared through centrifugation at 28 000 x *g* at 4°C for 40 mins followed by filtration of the soluble fraction through gauze. The cleared lysate was incubated with 1ml of GFP selector beads 50% slurry (Nanotag) pre-equilibrated in lysis buffer. Beads were incubated for 3 hours at 4°C under continuous rotation. Beads were washed with 60x slurry volume in lysis buffer and eluted through overnight incubation at 4°C with 160μl of Human Rhinovirus (HRV)-3C protease (Millipore) in a final volume of 2-3ml. Following elution, the eluate was collected through gentle centrifugation of the beads at 1000 x *g* and collection of the supernatant. The beads were then washed in double the eluate volume with wash buffer (50mM Tris HCl pH 8.0, 50mM (NH_4_)_2_SO_4_, 1mM MgCl_2_, 10μM ZnCl_2_, 10mM β-mercaptoethanol) and the resulting wash fraction combined with the eluate to dilute the (NH_4_)_2_SO_4_ to ∼120mM. The eluate mixture was further diluted through addition of an equivalent volume of Tris buffer (50mM Tris HCl, 1mM MgCl_2_, 10μM ZnCl_2_, 10mM β-mercaptoethanol) to reduce the final (NH_4_)_2_SO_4_ concentration to ∼60mM. Next, the eluate was loaded onto a MonoQ GL 5/50 column (GE Healthcare) and eluted in a linear gradient from 60mM to 1M (NH_4_)_2_SO_4_ in 50mM Tris HCl pH 8.0, 1mM MgCl_2_, 10μM ZnCl_2_, 10mM β-mercaptoethanol. MonoQ purification produced two peaks corresponding to RNA polymerase I (eluting at ∼380mM (NH_4_)_2_SO_4_) and RNA polymerase III (eluting at ∼550mM (NH_4_)_2_SO_4_). Human RNA polymerase III fractions were collected and diluted to a final (NH_4_)_2_SO_4_ concentration of ∼110mM. The sample was then concentrated using a Vivapsin 500 (100 000 MWCO) to a final concentration of approximately 0.05-0.1mg/ml. The concentrated sample was used immediately for grid preparation.

### RNA Elongation and Cleavage Assay

1,5 µl of human Pol III (∼0,1 mg/ml) was preincubated with 0.25 pmol of pre-annealed minimal nucleic acid scaffold (template DNA: 5’-CGAGGTCGAGCGTTGTCCTGGT-3’, non-template DNA: 5’-CGCTCGACCTCG-3’; RNA: 5’-FAM-AACGGAGACCAGGAC-3’) in transcription buffer (20 mM Hepes pH 7.8, 60 mM (NH_4_)_2_SO_4_, 8 mM MgSO_4_, 10 µM ZnCl_2_, 10% (v/v) glycerol, 10 mM DTT) for 20 min at 20°C. For RNA elongation, NTPs (1,4 mM end concentration each) were added and the reaction was incubated for 30 or 60 min respectively at 28°C. To examine cleavage activity, the preincubated reaction was incubated for 30 min at 28°C without the addition of NTPs. To stop the reaction an equal amount of 2x RNA loading dye (8 M Urea, 2x TBE, 0,02% bromophenol blue, 0,02% xylene cyanol) was added and the sample was heated to 95°C for 5 min. As control 0.25 pmol of scaffold was treated identically, without addition of polymerase and NTPs. 0.125 pmol of FAM-labeled RNA product (as well as a marker containing 9 nt, 15 nt and 21 nt long FAM-labeled RNAs: 5’-FAM-GACCAGGAC-3’, 5’-FAM-AACGGAGACCAGGAC-3’, 5’-FAM-UGUUCUUCUGGAAGUCCAGTT-3’) was separated by gel electrophoresis (20% polyacrylamide gel containing 7 M Urea) and visualized with a Typhoon FLA9500 (GE Healthcare).

### CryoEM Sample Preparation and Data Collection

Human Pol III cryoEM samples were prepared on C-Flat 1.2/1.3 (400 mesh) grids coated with a thin film of continuous carbon prepared in house. Grids were glow discharged at 15mA for 30s using a PELCO EasyGlow instrument prior to sample addition. A 3μl volume of sample at ∼0.06mg/ml concentration was applied and incubated for 30s at 18°C and 100% humidity. Grids were blotted for 1s at blot force 1 with a 0.5s drain time and plunge frozen in liquid ethane using the VitroBot Mark IV system (FEI).

Data collection was carried out using a FEI Titan Krios transmission electron microscope (Thermo Fisher) operating at 300KeV and equipped with a Falcon III direct electron detector (Astbury Biostructure Laboratory, University of Leeds). Separate data collections were carried out for both untilted and 30°tilted datasets. All datasets were imaged using EPU automated acquisition software with the Falcon III operating in electron counting mode at a nominal magnification of 75 000x and a calibrated sampling of 1.065 Å/pixel. For untilted data collection, 3115 movies were collected. Movies were collected over 45 frames with a 70s exposure time and a total dose of 44.1 e^-^, giving a dose per frame of 0.98 e^-^/Å^2^ and a dose rate of 0.63 e^-^/Å^2^/s. Data was collected over a defocus range of -1μm to -3μm. Tilted data collection was carried out at 30°in two separate sessions. The first session collected 921 movies, with a total dose of 37.8 e^-^ fractionated over 38 frames during a 70s exposure, yielding a dose per frame of 0.99 e^-^/Å^2^ and a dose rate of 0.54 e^-^/Å^2^/s. The second session collected 1703 movies, imaged with a total dose of 40.6e^-^ fractionated over 38 frames during a 70s exposure. This gave dose per frame of 1.07 e^-^/ Å^2^ and a dose rate of 0.58 e^-^/Å^2^/s. In both tilted data collections, micrographs were collected using a -1.2μm to -3μm defocus range.

### CryoEM Image Processing

Frame alignment and dose weighting was carried out on-the-fly using MotionCor2 ^39^. Following motion correction, CTFFIND4 implemented in the cisTEM software package was used for contrast transfer function (CTF) estimation ^40^. Particle picking was carried out using the *ab initio* particle picking option in cisTEM ^41^ and resulting particles exported to Relion 3.1 ^42^. Subsequent 2D and 3D classification, refinement and post-processing steps were carried out using Relion 3.1 and *ab inito* model generation using Cryosparc v2 ^43^. For the untilted dataset, 332 238 particles were selected, yielding a final particle set of 139 891 particles corresponding to hPol III following multiple rounds of 2D classification. This particle subset was used to generate an initial model of the hPol III structure using the *ab initio* model functionality in Cryosparc v2. Similarly, 87 075 particles were selected from 2624 30 tilted micrographs, yielding 32 787 particles following 2D classification. Both particle sets were combined generating the merged particle set of 172 678 particles. This was subject to 3D classification in Relion 3.1 using the Cryosparc *ab inito* model as a reference. Classification produced 5 classes, with a single class (class 4, containing 68 291 particles) corresponding to the complete polymerase molecule. This class was refined and subject to CTF refinement. This was a sequential procedure, first correcting for trefoil and 4^th^ order aberrations, followed by correction for magnification anisotropy in the second step. In the final step the defocus was refined on a per particle basis to correct for errors in CTF estimation for tilted particles. Following this, a further refinement was carried out, yielding a model at 3.7 Å resolution at the gold-standard 0.143 FSC cut-off criterion. Following refinement and post processing, the map was filtered according to the local resolution of each region using the local resolution functionality implemented in Relion 3.1. Inspection of the resulting density revealed poor density and low resolution of the heterotrimer region. In order to improve this region, a local mask was generated, and 3D classification carried out without alignment localised to the heterotrimer. This produced 3 classes, of which one (containing 25 369 particles) produced a model with improved heterotrimer density following consensus refinement. This was subject to further CTF refinement, consensus refinement and local resolution estimation to produce the final model reporting 4.0 Å global resolution at the 0.143 FSC cut-off criterion.

### CryoEM Model Building and Refinement

As an initial step, homology models were generated for all core (RPC1, RPC2, RPC10, RPAC1, RPAC2, RPABC1, RPABC2, RPABC3, RPABC4 and RPABC5), heterodimer (RPC4 and RPC5) and stalk (RPC8 and RPC9) subunits using the PHYRE2 webserver ^44^. These were rigidly fitted into the locally-filtered map using the fitted yeast apo RNA Pol III structure (RCSB Protein Data Bank (PDB) code: 6EU2) as a guide for the relative positioning of the subunits in UCSF Chimera ^45^. The placed homology models were then fitted manually to the density using the COOT software package ^46^, at this stage regions not present in the EM density were removed from the model. Following manual fitting, the model was fitted using the real-space refinement functionality in PHENIX ^47^. The lower resolution of the more dynamic heterotrimer region did not permit the use of this strategy for these subunits. Therefore, the existing crystal structure of human RPC3 in complex with a fragment of RPC7 ^48^ (RCSB PDB Code: 5AFQ) was structurally aligned with the yeast heterotrimer in the yeast apo-RNA Pol III structure in UCSF Chimera. Comparison revealed a highly similar structure and relative arrangement for the RPC3 and RPC7 regions present in both structures. Further to this, a homology model for RPC6 was generated using PHYRE2. This was structurally aligned in UCSF Chimera to the yeast RPC6 subunit present in the yeast apo Pol III structure. The region of human RPC6 consisting of residues 174-289 was selected for inclusion in the model, as this was the region which corresponded to residues 171-271 of yeast RPC6 which were visible in the yeast RNA Pol III apo state ^4^. Following selection of the relevant models, they were positioned using the yeast apo structure as a guide and then fitted to the human EM map using the sequential fit option in UCSF chimera.

### RPC5 Cloning, Expression and Purification

Based on secondary structure predictions using PsiPred ^49^ and HHPred programs ^50^ we designed 13 different constructs of the C-terminal extension of Rpc5 subunit (Uniprot ID Q9NVU0). A PCR-based strategy was used to amplify fragments of Rpc5-WH1/2, Rpc5-WH3/4 and Rpc5-WH1/4 from its genomic DNA (Genscript). The constructs were subsequently cloned into pOPINF or pOPINJ plasmids for bacterial expression or into pACEBac1 plasmid for baculovirus–insect cells expression. Two hexahistidine-tagged constructs that rendered high yield expression of undegraded proteins (Rpc5 (259-440) and Rpc5 (556-708)) were selected for large-scale production. Both protein constructs were expressed and purified following the same protocol. Cells were grown at 37°C, 200 rpm in Terrific Broth (TB) to OD_600_= 1.5 and protein expression was induced with 1mM IPTG at 20°C overnight. All subsequent steps were performed at 4°C. Harvested cells were resuspended in 20 mM HEPES pH 7.9, 150 mM NaCl, 10 mM imidazole and 10 mM β-mercaptoethanol supplemented with DNAse I and two protease-inhibitor tablets (Roche). After a 30-min incubation, the sample was sonicated and fractionated by centrifugation at 20 000 rpm for 40 min. Then, the soluble fraction was loaded in a HisTrap HP 5 mL affinity column (GE Healthcare) pre-equilibrated with lysis buffer. After extensive washes of the chromatographic column, the protein was eluted with lysis buffer supplemented with 250 mM imidazole. The sample was diluted to 70 mM NaCl and injected into an HiTrap Heparin HP 5 ml (GE Healthcare) column equilibrated with 20 mM HEPES pH 7.9, 70 mM NaCl and 10 mM β-mercaptoethanol. The protein was subsequently eluted using an isocratic gradient from 70 mM to 2 M NaCl in 30 column volumes (CV). Fractions containing Rpc5 constructs were identified by SDS-PAGE analysis. Cleavage of the His-tag was performed overnight incubating the protein with 3C protease in a 1:50 molar ratio (3C protease: Rpc5). Uncleaved His-tagged proteins were removed by incubation of the sample with 1 ml HisPur™ (Thermo Fisher) nickel resin for 1h at 4°C. The cleaved protein was concentrated to 5 ml and loaded in a HiLoad 16/600 Superdex 75pg gel-filtration column (GE Healthcare) equilibrated with 50 mM Tris-HCl pH 7.5, 150 mM NaCl and 10 mM β-mercaptoethanol. Purified Rpc5 (259-440) and Rpc5 (556-708) were concentrated to 30 mg/ml and 80 mg/ml, respectively flash-frozen and stored at -80°C.

The expression of selenomethionine-derivatized proteins was performed in the methionine-auxotroph *E. coli* B834(DE3) strain (Novagen) using SelenoMet™ medium (Molecular Dimensions) supplemented with SelenoMet Nutrient Mix and 40 mg/l L-selenomethionine (SeMet). Purification of the proteins was performed as described for the native proteins.

The expression of the whole C-terminal extension of Rpc5 (referred as Rpc5-WH1/4) was performed in the insect cells/baculovirus expression system. Large-scale suspension cultures (300 mL) of High Five insect cells at 0.5 × 10^6^ cells/ml were grown in Insect-Xpress media (Lonza) and inoculated with P2 baculovirus solution containing the Rpc5 constructs. Proliferation arrest was assessed by measurement of GFP production until fluorescence reached a plateau. Cells were harvested at 800 *x g* for 5 min and the pellets were stored at -20°C. After a milder sonication step, purification was performed following the protocol described above. Finally, protein was concentrated to 10 mg/mL, flash-frozen in liquid nitrogen and stored at -80°C.

### Crystallization, Data collection and Structure Determination

Crystals used for structure determination were grown from a 1:1 ratio solution (protein: reservoir) using the vapour diffusion technique at 4°C. Rpc5 (259-440) crystals in P6_1_22 space group were obtained at 30 mg/mL after 3-4 days equilibration in 3.2 M NaCl, 100 mM Sodium Acetate pH 4.6 and 10 mM ZnCl_2_. SeMet-Rpc5 (259-440) crystals in the same space group were obtained at similar conditions but required the use of streak seeding with diluted native crystals to favour nucleation. Rpc5 (556-708) crystals in P2_1_ space group grew at 35 mg/mL in 3.2-3.8 M Ammonium Acetate and 100 mM Bis-Tris Propane pH 6.5-7 after 2 weeks. Crystallization of SeMet-Rpc5 (556-708) protein was performed under identical conditions. All crystals were flash-frozen in liquid nitrogen using perfluoropolyether oil (Hampton Research) as cryoprotectant. A dataset from native Rpc5 (259-440) was collected at 0.9198Å-wavelength in I24 beamline of Diamond Light Source (DLS). Additionally, multi-wavelength anomalous dispersion (MAD) data collections were performed at the peak, remote and inflection wavelengths from SeMet-derivatized crystals of Rpc5 (259-440) and Rpc5 (556-708) in I03 beamline of DLS. Using the MAD dataset, an initial model of SeMet-Rpc5 (259-440) at 2.7 Å was determined with the SHELXC/D/E suite from the HKL2Map program (for phase determination) and the Buccaneer software (for model building). The native structure at 2.2 Å was solved by molecular replacement using the initial SeMet model as a search reference in PHENIX.automr. Subsequent refinement was performed using COOT and PHENIX suites. The structure of Rpc5 (556-708) at 1.48 Å was solved from the SeMet datasets using HKL2Map and Buccaneer programs and further refined to acceptable R_free_/R_work_ values with COOT and PHENIX. Protein secondary structure assignment from the atomic coordinates was performed using STRIDE ^51^.

### RPC5 SAXS Data Collection and Processing

SAXS data collection was carried out at the SWING small and wide-angle scattering beamline, SOLEIL Synchrotron, Saint Aubin, France. Purified RPC5 WH1/4 (164μM), WH1-2 (922μM) and WH3-4 (3691μM) were passed through a Bio SEC-3 HPLC column (Agilent) at 0.2ml/min in 50mM Tris-HCl pH 7.5, 150mM NaCl, 10mM β-mercaptoethanol with protein elution monitored using A_280_. Data were collected using an Eiger X 4M detector (Dectris) at a 2m distance, using a q range of 0 < q < 0.68Å^-1^. Data were reduced and buffer subtraction performed at the beamline. Data analysis was carried out using the ScÅtter software package for determination of radius of gyration (R_g_), P(r) distribution, particle maximum dimension (D_max_) parameters and for qualitative flexibility analysis (through generation of R_g_-Normalized Kratky, SIBLYS and Porod-Debeye plots) ^52^. Volumetric bead modelling was performed using the DAMMIN software package ^53^. Briefly, *ab inito* bead models were calculated using DAMMIN 10 times for each construct, fitting over the q range of 0 < q < 0.25 Å^-1^. The resulting bead models were averaged and filtered using the DAMAVER package ^54^, generating the final bead model reconstruction.

Comparison of the theoretical scatter profiles of the determined crystal structures for the WH 1-2 and WH 3-4 constructs was performed using the CRYSOL package for structural validation ^55^. Modelling of the entire RPC5 C-terminus (RPC5 WH1/4) was performed using Ensemble Optimisation (EOM) analysis from the ATSAS package ^56^ of the RPC5 WH1/4 SAXS data using the q range 0 < q < 0.2 Å^-1^. The determined WH1-2 and WH3-4 crystal structures were defined as rigid bodies in the RPC5 C-terminal sequence, with EOM analysis modelling the intervening 115 amino acid linker as dummy atoms, generating a pool of 10 000 random structural conformations from which the ensemble was selected to sample the continuous structural heterogeneity.

### Cell Culture

HEK293T cells (a kind gift from Dr Sebastien Guettler) were cultured in DMEM supplemented with 10% heat inactivated FBS and 1% penicillin/streptomycin at 37°C in 5% CO_2_. For transient knockdown, 20nM siRNA of either ONTARGETplus siRNA for RPC5 (Horizon Discovery) or AllStars negative control siRNA (Qiagen) was used per 6 well. This was transfected using Lullaby transfection reagent (Oz Biosciences) as per the manufacturers instructions. The sequences for each RPC5 siRNA are as follows: UGGAUAAGGCUGACGCCAA, GGGAGCAGAUUGCGCUGAA, CGACGAGACCAGCACGUAU, CCUCGAUGACCUACGAUGA. For transient overexpression of RPC5 (HA-tagged full length or ΔC), 1.5μg DNA was transfected in using Fugene HD transfection reagent (Promega) as per the manufacturers instructions.

### Antibodies

The following primary antibodies were used: POLR3A (ab96328, Abcam), POLR3B (ab137030, Abcam), POLR3D (ab86786, Abcam), POLR3E (ab134560, Abcam), HA tag (ab9110, Abcam), GAPDH (MAB374, diluted 1:5000 for Western Blotting, Merck) and GFP (ab290, Abcam). The following secondary antibodies were used: anti-rabbit IgG (H+L) DyLight™ 800 4x PEG conjugate (#5151, Cell Signalling Technology), anti-mouse IgG (H+L) DyLight™ 680 (#5470, Cell Signalling Technology). All antibodies were used at a dilution of 1:1000 unless otherwise stated.

### Co-immunoprecipitation and western blotting

HEK293T cells were seeded into 10cm plates in the presence of siRNA. After 48 hours they were subsequently transfected with pcDNA3.1+ N-HA carrying RPC5 (HA-tagged full length or ΔC) and maintained for a further 24 hours. Cells were lysed in RIPA buffer and co-immunoprecipitation was performed using Pierce™ anti-HA magnetic beads (ThermoFisher Scientific). Lysates were diluted in NuPAGE™ LDS 4x sample buffer (ThermoFisher Scientific) plus NuPAGE™ 10x sample reducing agent (ThermoFisher Scientific) before being boiled for 5 minutes. SDS PAGE was subsequently performed on the lysates in 4-12% Bis-Tris protein gels, transferred to nitrocellulose membrane, blocked for 1 hour in 5% milk/TBS/0.1% tween 20 and probed with primary antibody overnight at 4°C. Secondary antibodies were incubated for 1 hour at room temperature in the dark and detected using the Odyssey-CLx fluorescence imaging system (LI-COR Biosciences).

### Cycloheximide Chase Assay

HEK293T cells were seeded into 6 well plates in the presence of siRNA. After 24 hours they were subsequently transfected with RPC5 (HA-tagged full length or ΔC) and maintained for a further 24 hours. Cycloheximide was then added at a concentration of 300μg/ml and cells lysed at regular time points (0-8 hours) using RIPA buffer. Lysates were subsequently analysed via western blot, as previously stated.

## DATA AVAILABILITY

The electron density reconstructions and final model were deposited with the Electron Microscopy Data Base under accession code nos. EMD-xxx, and with the Protein Data Bank (PDB) under accession code xxxx.

The PDB accession numbers for the atomic coordinates and structure factors of the RPC5EXT tWHD1 and tWHD2 crystal structures reported in this paper are xxxx, and xxx, respectively.

## ACKNOWLEDGEMENTS

We thank all members of the Vannini and Engel groups for discussion throughout this study. We thank Tino Pleiner for discussion, Ilya Komarov for help with nuclear fractionation, Andrea Bleckmann and Juan Rodriguez Alcázar for assistance with confocal microscopy, and Astrid Bruckmann for mass spectrometry. We also thank Rebecca Thompson and the Astbury Biostructure Laboratory, University of Leeds for human Pol III imaging. In addition, we thank Ruth Peat and the Cell Services Scientific Technical Platform at the Francis Crick Institute for large-scale POLR1C-GFP cell growth. CE acknowledges funding by DFG’s Emmy-Noether Programme (EN 1204/1-1) and support by the collaborative research center 960 (SFB960/TP-A8).This work was funded by the Cancer Research UK Programme Foundation (CR-UK C47547/A21536) and a Wellcome Trust Investigator Award (200818/Z/16/Z) to A.V.

## AUTHOR CONTRIBUTIONS

C.E and P.G designed and performed genome editing experiments. P.G. created the stable POLR1C-sfGFP HeLa cell line. C.E., M.P. and J.L.D performed initial small-scale purifications of human Pol III. J.L.D optimized suspension cultivation of RPAC1-GFP cells, carried out transcription assays and confocal microscopy. G.A.-P. and E.P.R. performed large-scale purification of human Pol III and prepared samples for EM analysis and collected cryo-EM data. E.P.R. performed cryo-EM data processing. G.A.-P. performed x-ray crystallography experiments and analysed the data. J.G collected SAXS data. E.P.R and G.-A.P. analysed SAXS data. H.K. performed experiments in living cells. G.A.-P., E.P.R., H.K. and A.V. interpreted the data. A.V wrote the manuscript with input from all the authors. C.E and A.V. designed and supervised research.

---

**Extended Data Figure 2.**
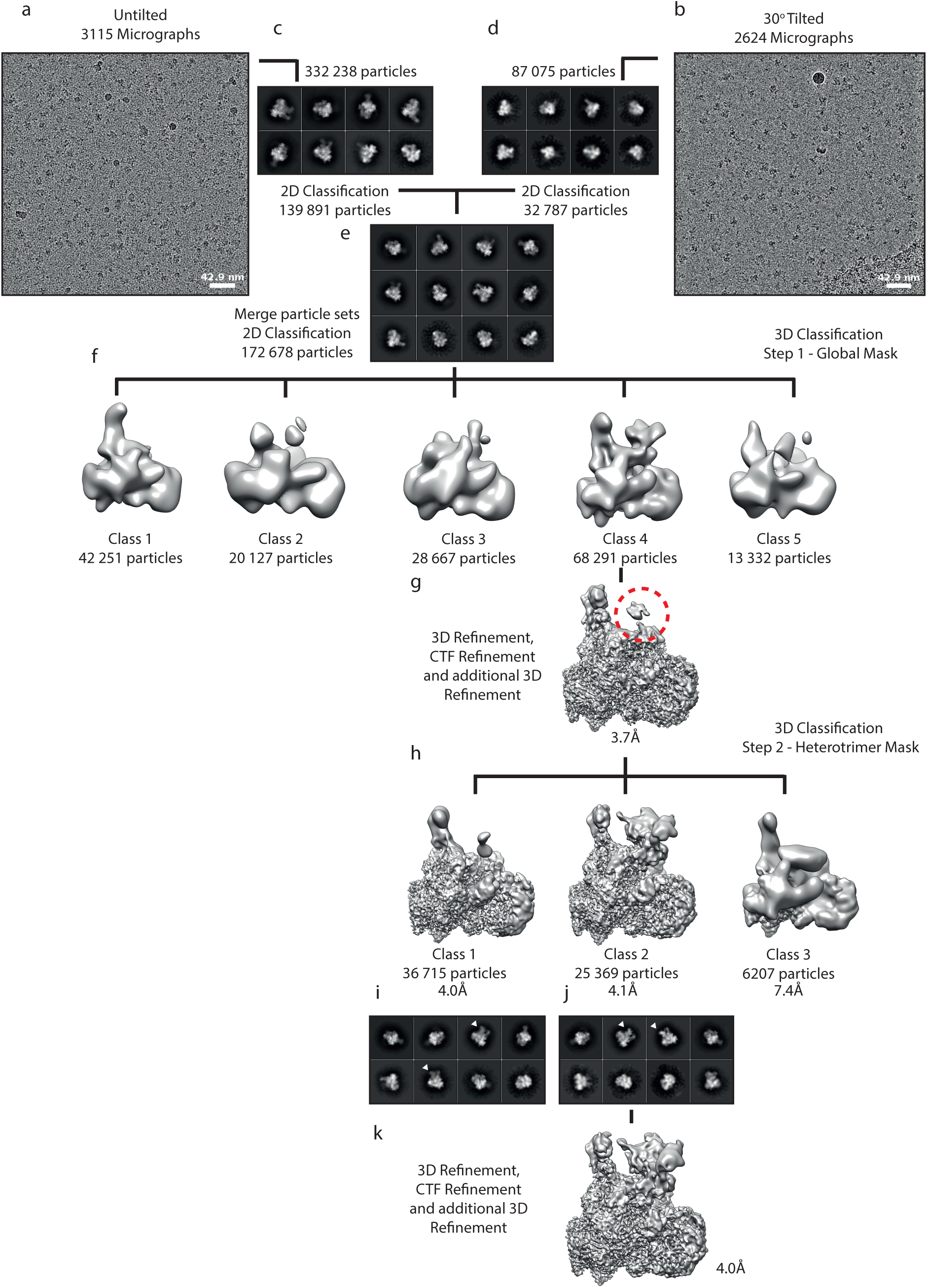
Human Pol III cryo-EM data processing. (a) Representative micrographs for untilted and (b) 30° tilted datasets Representative 2D class averages for (c) untilted, (d) 30° tilted and (e) merged datasets. (f) Consensus 3D classification of the resulting particle set using a cryosparc ab inito model as reference. The class corresponding to full polymerase was refined (g) and then subject to masked classification around the heterotrimer (marked with red circle) (h). (i,j) Representative 2D classes of resulting classes, with the heterotrimer density marked. The class corresponding to the full polymerase molecule was subsequently refined to generate the final model (k).

**Extended Data Figure 3.**
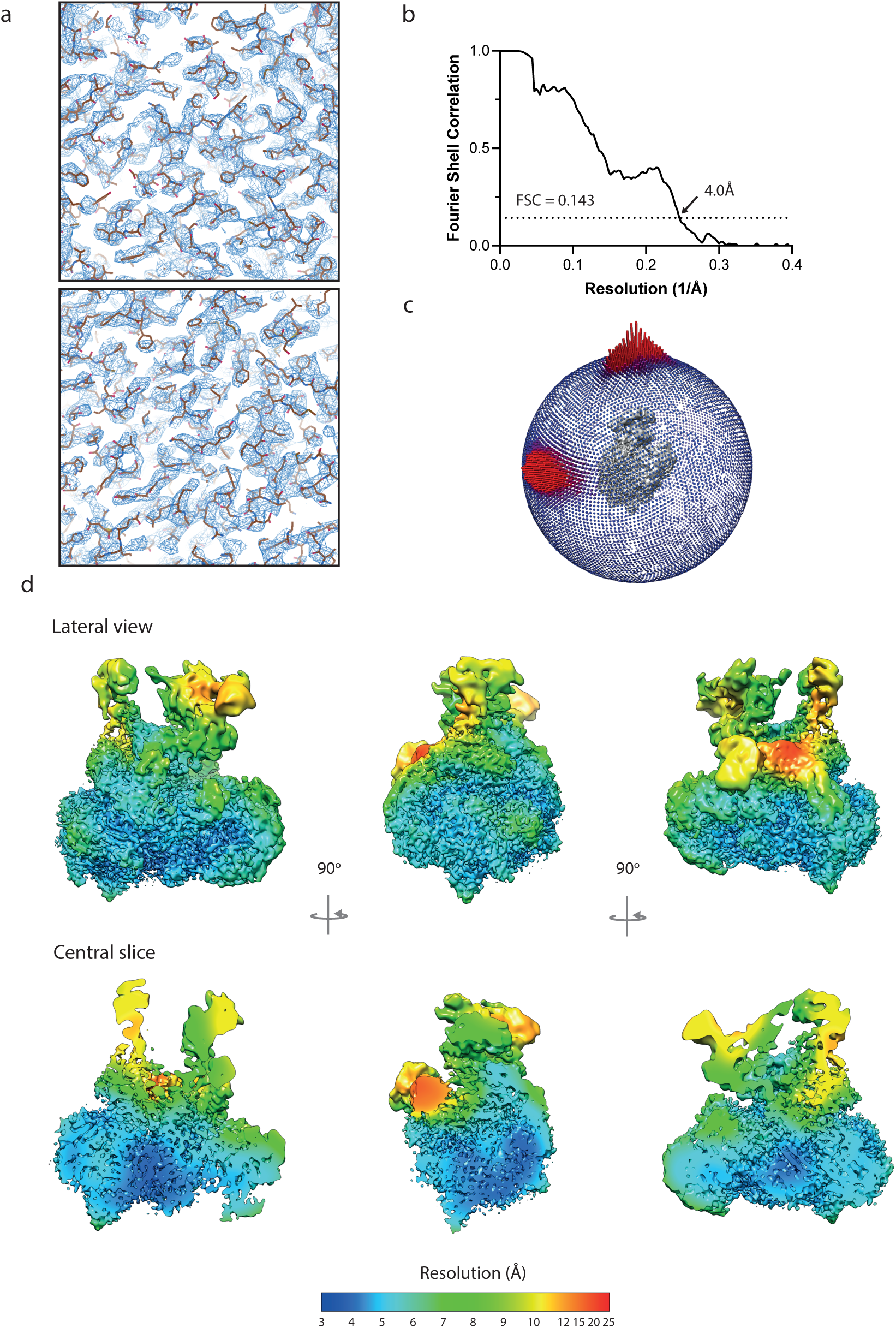
Resolution determination of human Pol III cryo-EM reconstruction. (a) Central slices of the refined map with model fitted. (b) Fourier shell correlation (FSC) of the final human polymerase III reconstruction, reporting a resolution of 4.0Å at 0.143 FSC. (c) Lateral view of the orientation distribution of the particles which contributed to the reconstruction. The heights of the bars reflect the relative number of particles in each orientation. (d) Lateral view (above) and central slice (below) displaying the local resolution distribution of the final refined map. Map density is filtered and coloured according to the local resolution estimation, as indicated by the colour key (below). Local resolution was estimated using the Relion-3.1 package.

**Extended Data Figure 4.**
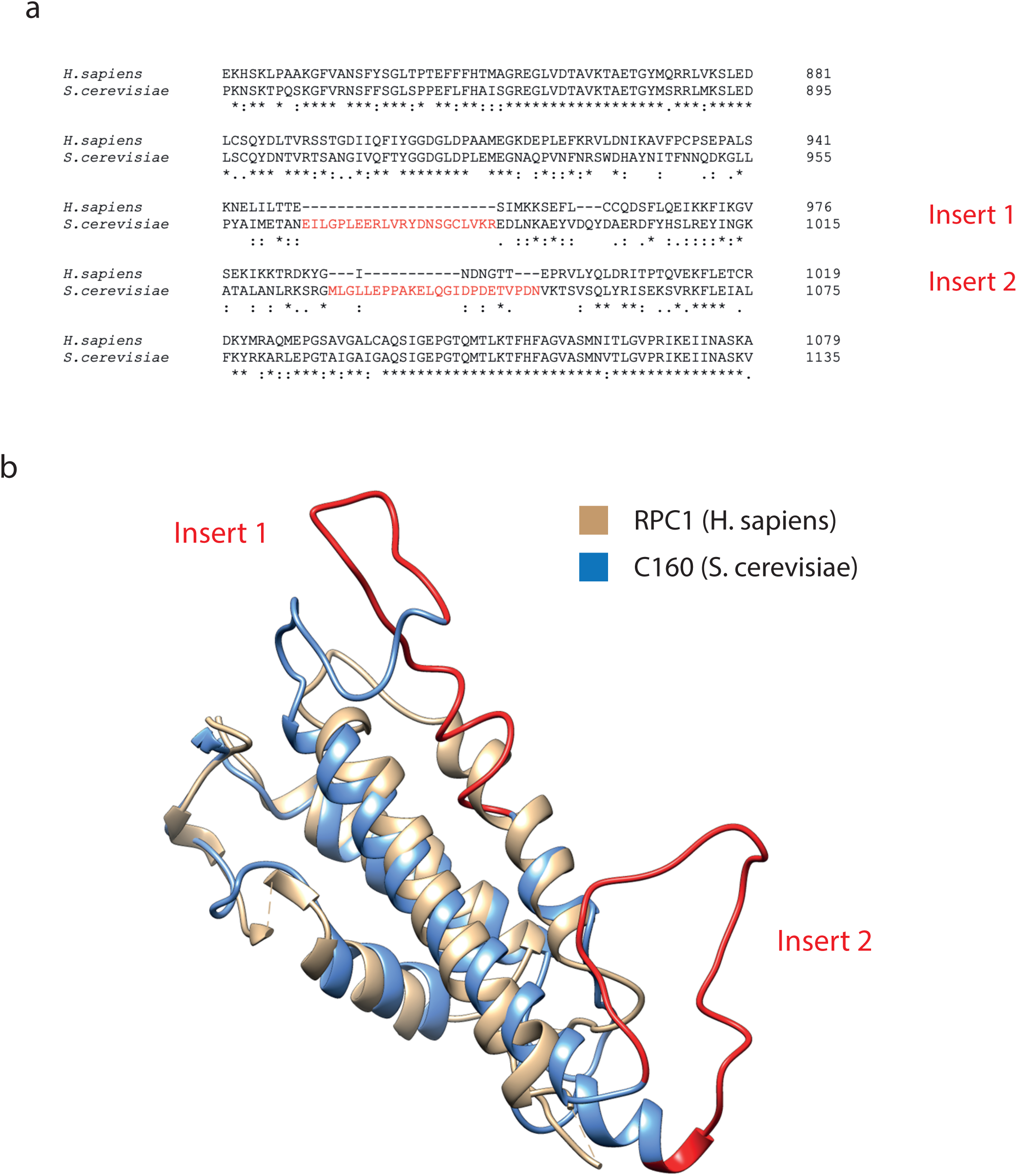
Comparison of the human and yeast foot domain in RPC1. (a) Sequence alignment of the foot region present in RPC1 and C160 showing two insertions in the yeast sequence relative to the human subunit, labelled insert 1 and 2. (b) Structural alignment of the human and yeast foot region showing a truncated structure for the human relative to the yeast homologue. Highlighted in red are the two insetions in the C160 sequence.

**Extended Data Figure 5.**
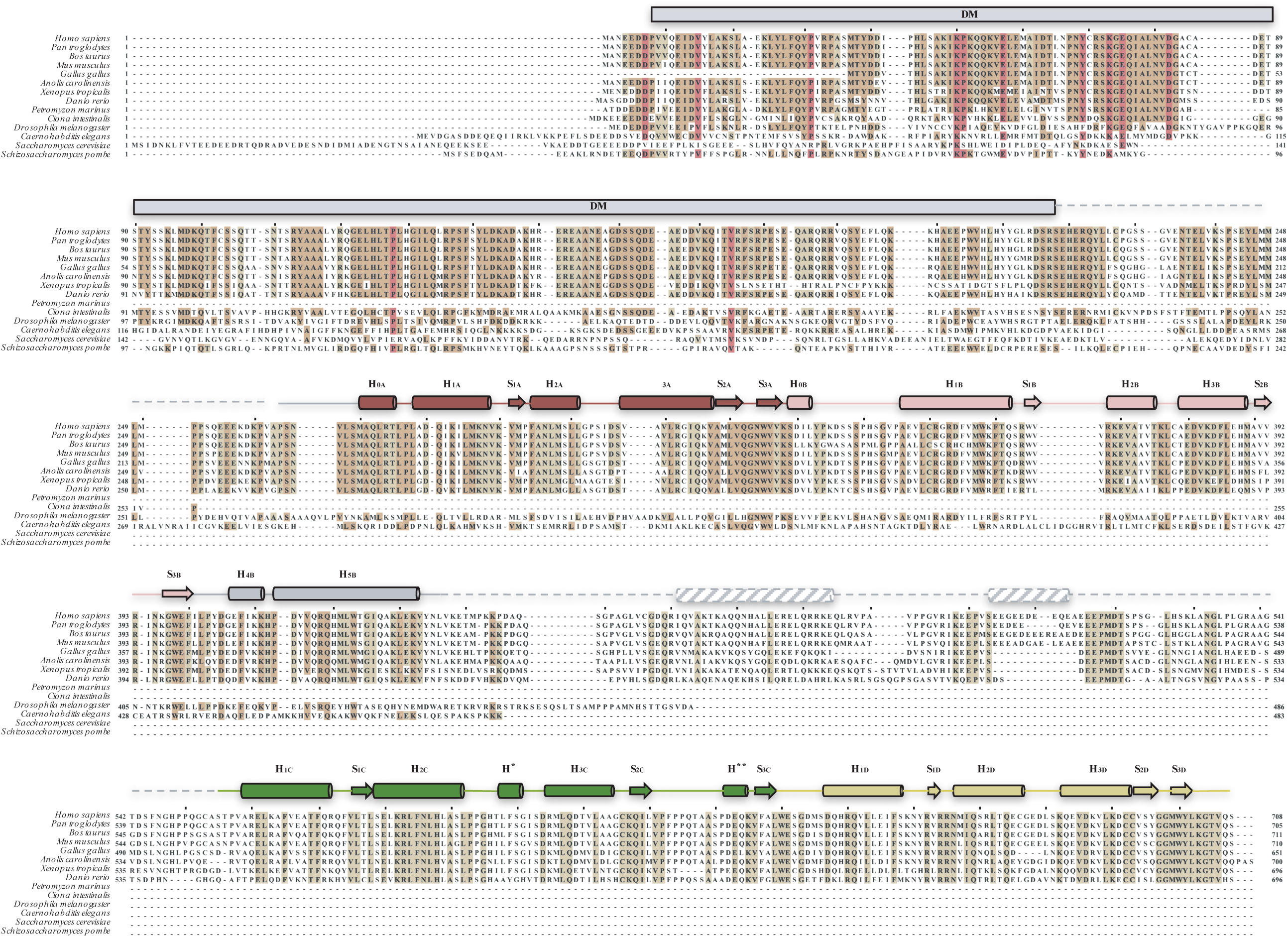
Rpc5 sequence and architecture conservation. Multiple sequence alignment (MSA) of human Rpc5 with representative organisms. Residues are coloured according to the identity percentage: >80% (red), >60% (dark brown) or >40% (light brown). A schematic representation of the known and predicted structural elements is depicted above the MSA. Solved structures of the linkers (grey), Rpc5-tWHD1 (red) and Rpc5-tWHD2 (green) are shown as cylinders (α-helixes) or arrows (β-strands). The secondary structure prediction of Rpc5 linker (between tWH1 and tWH2) is shown in dashed grey. Rpc5 dimerization module is shown as a grey box.

**Extended Data Figure 6.**
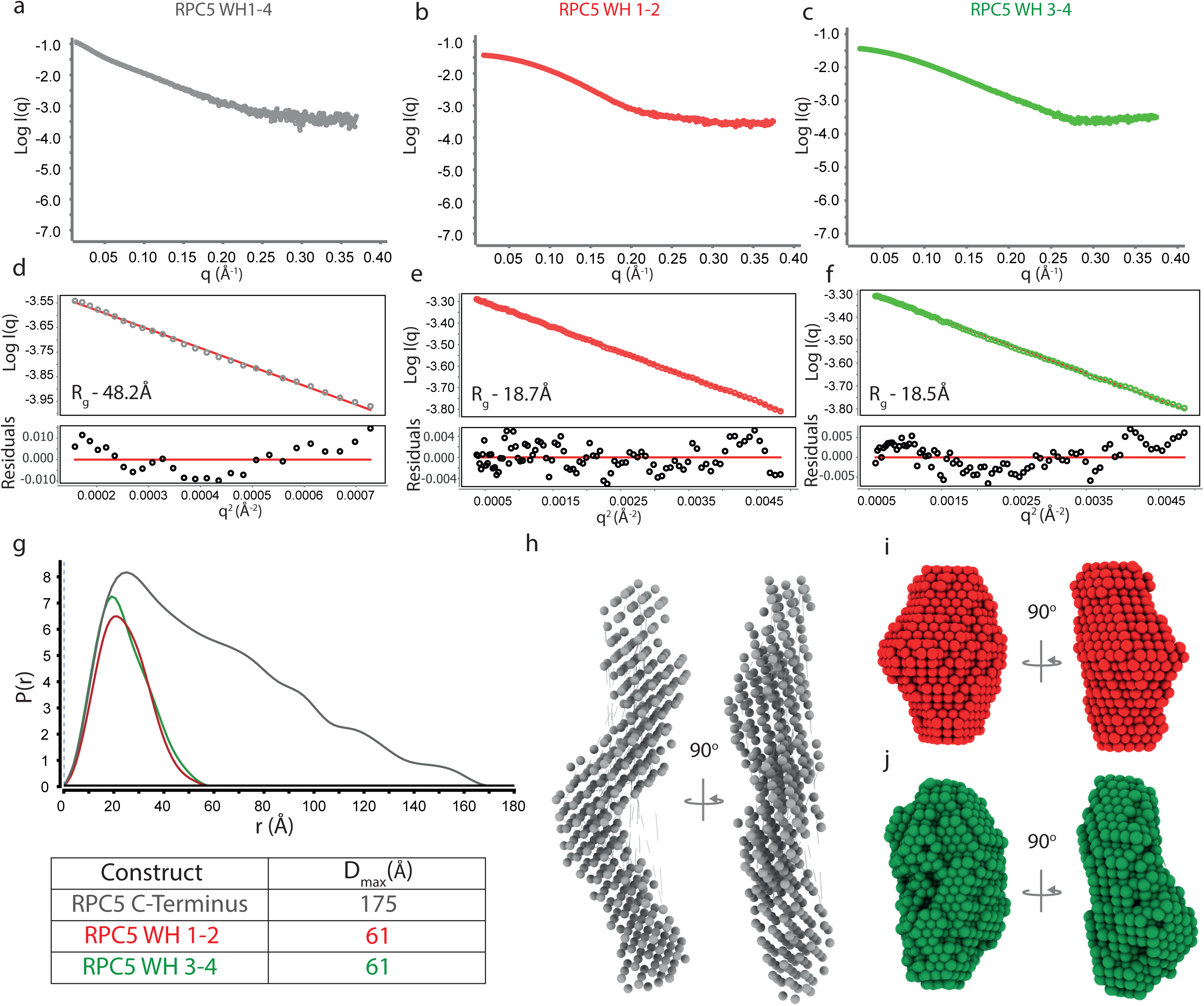
SAXS analysis of the RPC5 C-terminus. (a,b,c) One-dimensional scatter curve recorded for the (a) RPC5 WH1-4, (b) RPC5 WH1-2 and (c) RPC5 WH 3-4. (d,e,f) Guinier fitting for the recorded scatter curve. Shown is the fitting with the calculated R_g_ values inset (above) and the residuals for the Guinier fitting across the q-range of the Guinier region (below) for (d) RPC5 WH1-4, (e) RPC5 WH 1-2 and (f) RPC5 WH3-4. (g) Overlayed P(r) distruibutions for all constructs (above) with the particle maximum diameter (D_max_) values determined at the x-intercept (below). Averaged DAMMIN dummy-atom bead models calculated for (h) RPC5 WH1-4, (i) RPC5 WH 1-2 and (j) RPC5 3-4 constructs.

**EXtended Data Figure 7.**
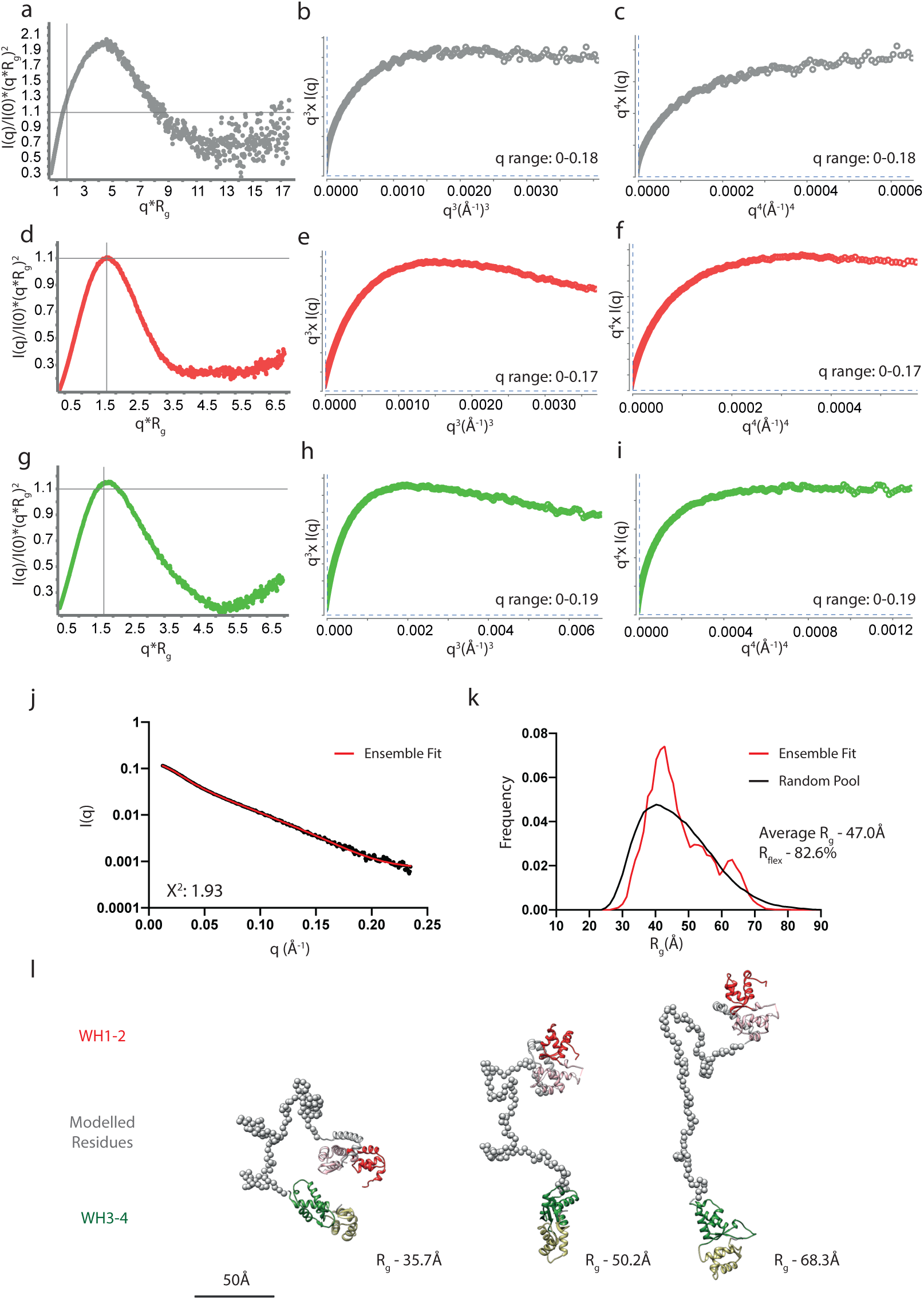
Qualitative assessment of protein flexibility from SAXS data. Calculated R_g_ normalized kratky plots for (a) RPC5 WH1-4, (d) RPC5 WH 1-2 and (g) RPC5 WH 3-4. All plots display a bell-shaped profile, indicative of folded protein. Shown also is the intersection at 3^½^, 1.1 (crosshairs). A peak observed at this point reports a globular structure, as observed in (d) and (g). Dispersion of the peak away from this point in (a) suggests a more elongated structure. Calculated SIBLYS plots for RPC5 WH1-4 (b), RPC5 WH1-2 (e) and RPC5 WH3-4 (h). These are comapred to Porod-Debye plots shown for RPC5 WH1-4 (c), RPC5 WH1-2 (f) and RPC5 WH3-4 (i) over the same q-range in each pairwise comparison. The RPC5 WH1-4 shows a plateau in the SIBLYS plot before the Porod-Debye, suggestive of a flexible protein. Both RPC5 WH1-2 and RPC5 WH3-4 show a prominent plateau in the Porod-Debye, suggestive of a more rigid body. This suggests the RPC5 C-terminus consists of two rigid bodies with a flexible linker between. (j,k,l) Ensemble optimisation (EOM) analysis of RPC 5 C-terminus. Both winged-helix crystal structures were defined as rigid bodies, with EOM modelling the additional 115 residue linker in the RPC5 sequence as dummy atoms. (j) Comparison of the theoretical scattering of the selected ensemble to the experimental SAXS curve for RPC5 WH1/4, with inset the reported fit X^2^ value of 1.93 over the q range 0<q<0.2Å^-1^. (k) Comparison of the R_g_ distribution of the random pool of 10000 models (black) to the selected ensemble (red). The reulting ensemble reported an average R_g_ of 47.0Å and an R_flex_ of 82%. The selected ensemble displayed a wide dispersion of R_g_ values, equivalent to the range of the random pool, suggesting together with the high R_flex_ value a flexible structure. (l) Selected models from the fitted ensemble with the reported R_g_ for each showing a range of sizes modelled by the EOM analysis.

**Extended Data Figure 8:**
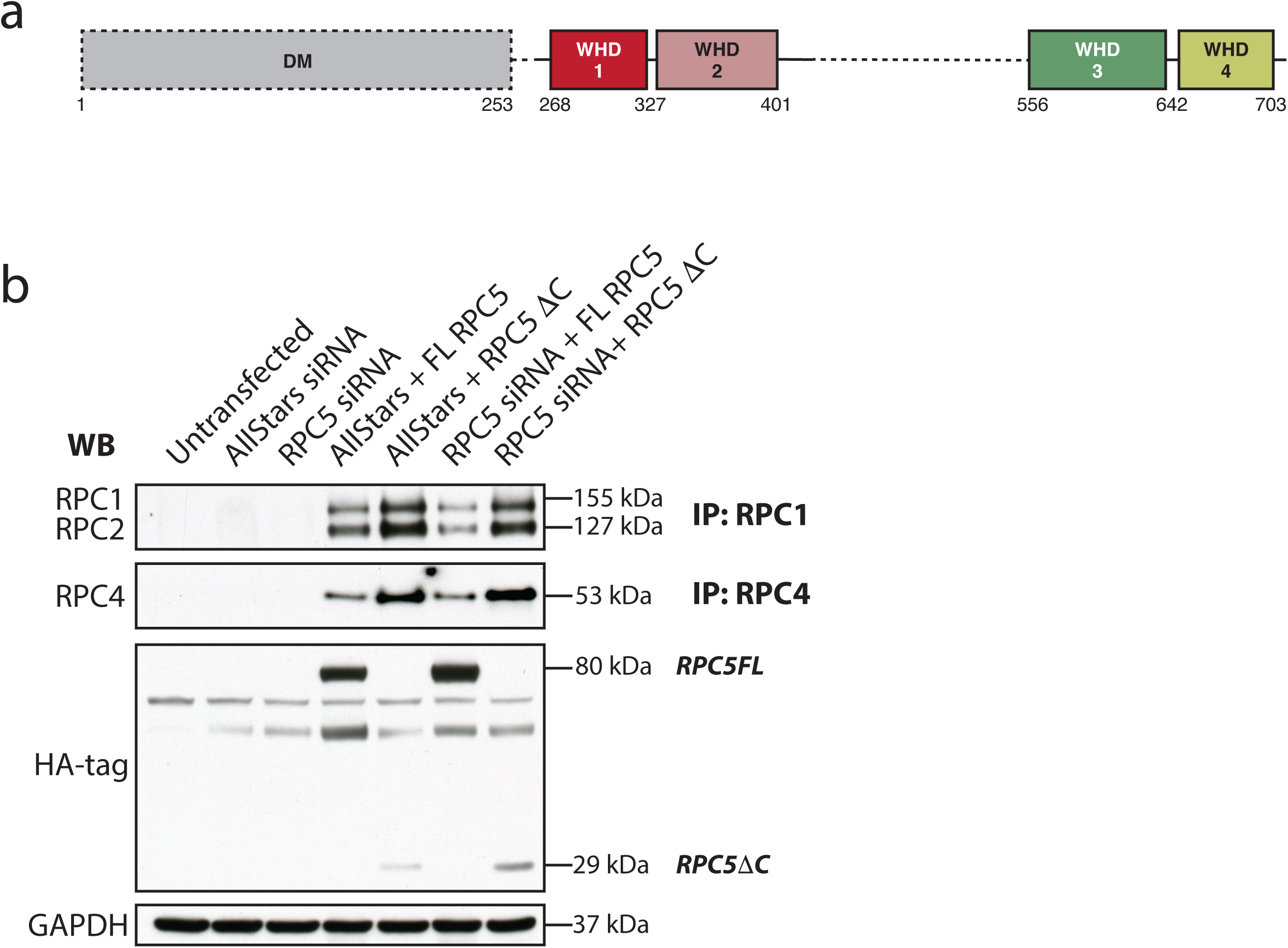
The domain structure of RPC5 is shown (a). N-terminally, HA-tagged DNA constructs were ordered constituting either amino acids 1-708 RPC5 Full Length (FL) or amino acids 1-253 (DC) where the tandem wing helix domains had been removed. Coimmunoprecipitation in HEK293T cells were performed to investigate whether truncated RPC5 would affect RNA Pol III complex assembly (b). Endogenous RPC5 was knocked down using siRNA before either FL or DC HA-tagged RPC5 constructs were transfected in. Magnetic HA-beads were used to pull down components of the RNA Pol III complex (RPC1, 2 and 4). From this is was concluded that truncation of RPC5 does not inhibit Pol III complex assembly

**Extended Data Figure 9.**
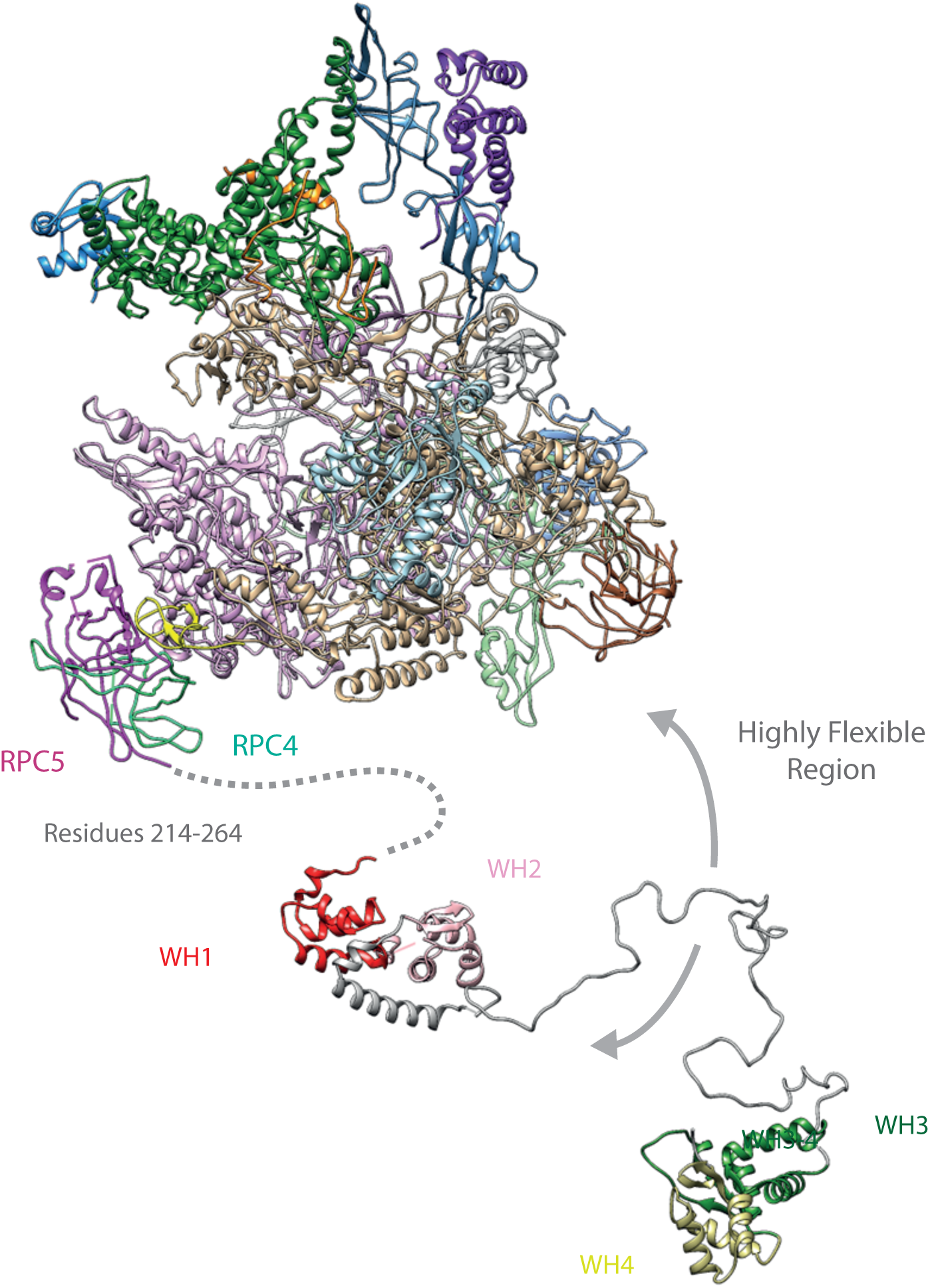
Superimposition of human RNA polymerase III structure and modelled RPC5 C-terminus, distinct regions of RPC5 are labelled.

**Extended Data Table 1.**
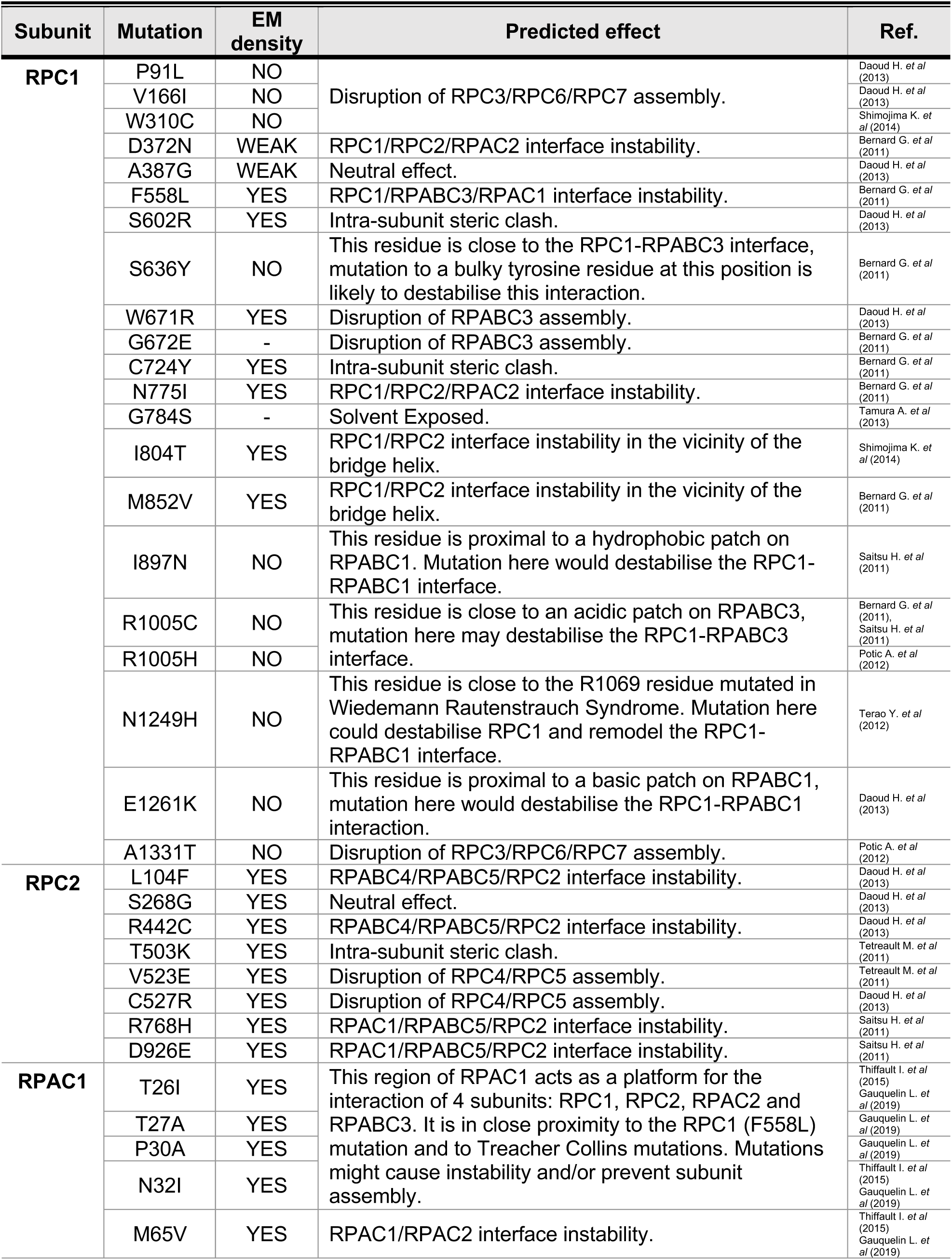
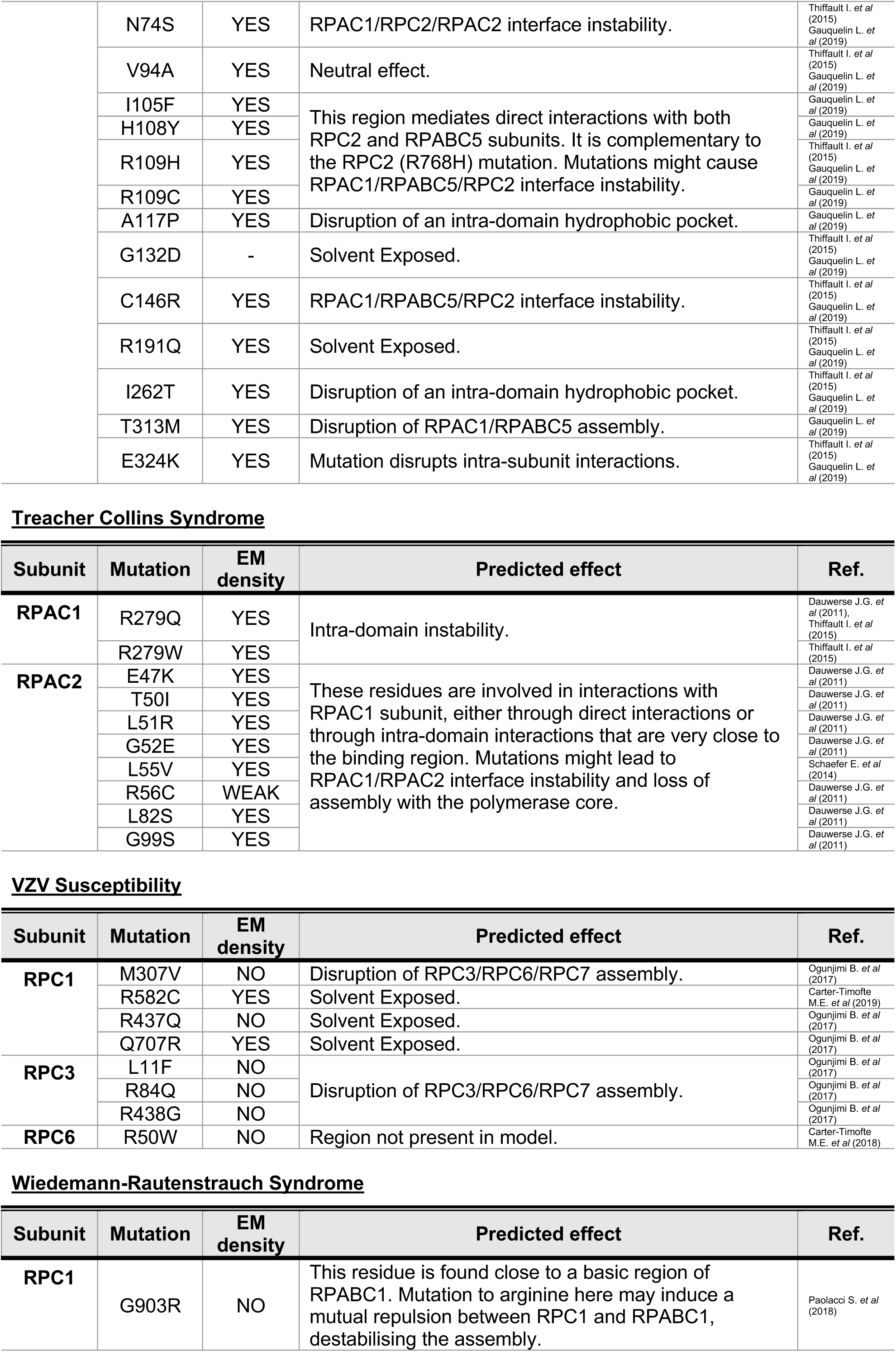
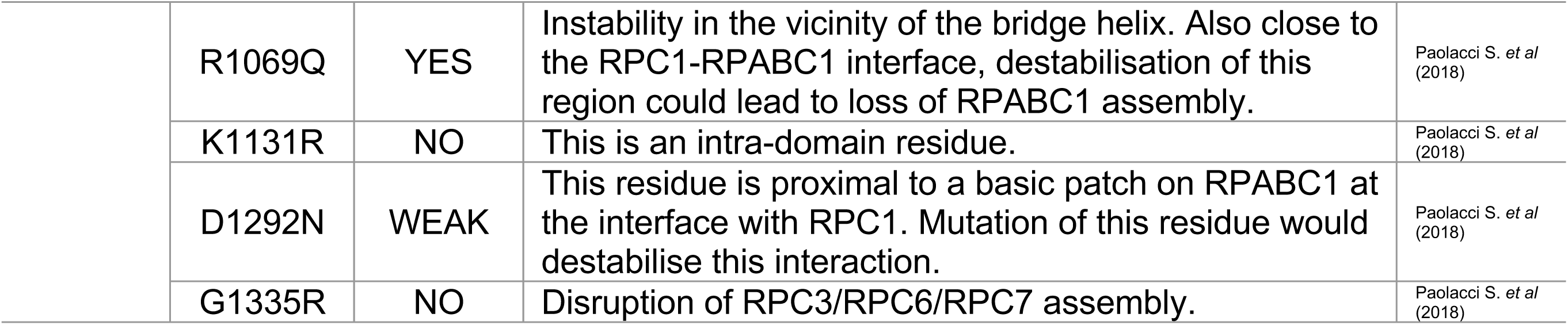
Overview of human RNA Polymerase III mutations. Hypomyelinating Leukodystrophy

